# Integrated stress response activation in combination PG3 and cisplatin-treated *TP53*-mutated head and neck squamous carcinoma cells induces potent apoptotic response

**DOI:** 10.64898/2026.09.22.752907

**Authors:** Xiaobing Tian, Praveen R. Srinivasan, Wafik S. El-Deiry

## Abstract

Cisplatin remains as standard chemotherapy for patients with HNSCC, but rapid development of drug resistance has limited patient benefit. The p53 tumor suppressor plays a central role in the cellular response to DNA damage in cancer, triggering apoptosis to prevent propagation of damaged cells in tumor development. *TP53* gene mutations occur in 65-86% of HNSCC. Small molecule PG3 induces the integrated stress response (ISR), leading to apoptosis via the HRI-eIF2α-ATF4-PUMA axis. We hypothesized that a combination of PG3 plus cisplatin could increase apoptosis in *TP53*-mutated HNSCC cells through enhanced induction of the ISR and ATF4. PG3 synergized with cisplatin to inhibit cell viability, leading to potent apoptosis in *TP53*-deficient cells. The effect was regulated through the HRI-ATF4-NOXA pathway. Furthermore, we identified that cisplatin activates HRI and leads to the degradation of CReP (constitutive repressor of eIF2α phosphorylation) via E3 ligase β-TrCP, contributing to the induction of the ISR. We noted decreased ATF4 levels after treatment with cisplatin, CPT, or PG3 and cisplatin. Thus, combined therapy of PG3 plus cisplatin likely results in adaptation and acquired resistance via degradation of ATF4. We targeted the degradation mechanism of ATF4 by inhibiting β-TrCP1, CK1δ, or CK2, respectively. Each approach successfully blocked ATF4 degradation induced by cisplatin or PG3 plus cisplatin and enhanced apoptosis. Our results provide a rational strategy for triple treatments, involving an ISR inducer, a DNA damaging drug, and a β-TrCP inhibitor/CK1δ inhibitor/CK2 inhibitor, to achieve potent and prolonged anti-tumor effects and overcome chemoresistance in HNSCC.

## Introduction

Head & neck squamous cell carcinoma (HNSCC) occurs in the mucosal epithelial tissue of the oral cavity, oropharynx, and larynx. About 60% of stage III or IV HNSCC patients are treated with chemotherapy (1). Cisplatin is a standard chemotherapy drug for stage III and IV HNSCC patients. In addition, the mutation frequency of the *TP53* gene in HNSCC is 65–85% (2). Cisplatin molecules enter cells through the chloride transport receptor 1 (CTR1). A cisplatin molecule is activated by monoaquation and diaquation in cells through a nucleophilic substitution reaction. Then, activated cisplatin molecules form covalent bonds with DNA. Most cisplatin DNA adducts are intra-strand DNA species (65%) and inter-strand crosslinks (25%) (3). Multiple cisplatin-resistance mechanisms have been reported (4). Adaptation to ER stress in HNSCC OECM1 cells leads to a decrease of CHOP (C/EBP homologous protein) and upregulation of DNA polymerase η, leading to resistance to cisplatin through trans-lesion DNA repair (5). Recently, it was demonstrated that ATF4 (Activating transcriptional factor) downregulation promoted cisplatin resistance in p53-mutated SGC7901 and p53-deleted BGC823 gastric cancer cells (6).

The integrated stress response (ISR) is a conserved signaling pathway that helps cells adapt to various stresses. ISR is mediated by four kinases, PERK, GCN2, PKR, and HRI, via phosphorylation of eIF2α at Ser51 in eukaryotic cells. Amino acid deficiency activates GCN2, ER-stress activates PERK, double-stranded RNA virus infection activates PKR, while heme deficiency activates HRI (7, 8). eIF2α phosphorylation causes inhibition of 5’-cap-dependent global translation, which triggers the ISR. ATF4 is selectively translated due to the presence of 2 upstream open reading frames (uORFs) in its 5’-untranslated region (5’-UTR) of mRNA, which are bypassed by ribosomes under stress conditions, allowing efficient translation of the ATF4 coding region. ATF4 is a basic region-leucine zipper transcriptional factor and binds to promoter regions of its target genes, specifically C/EBP-ATF Response Element (CARE) sequences and regulates transcription (7–9). ATF4 is a master effector of the ISR and regulates pro-survival or pro-apoptotic pathways. Compared to normal cells, due to faster growth and proliferation, cancer cells usually have proportionally elevated ISR to protect them. When a small molecule can trigger ISR, and result in too intense or prolonged stress, this will shift the balance from protection to apoptosis (7, 8, 10, 11).

DNA damage triggers the ISR and selective translation of cell cycle regulator p21(WAF1), which is dependent on the uORFs in the 5’-UTR of their mRNAs (12, 13). DNA damage following chemotherapy treatments can activate apoptosis via p53 in *TP53* wild-type cancer cells. In *TP53*-mutated/deleted cancer cells, DNA-damaging drugs induce cellular apoptosis through other mechanisms, such as the integrated stress response (14). Sharma *et al*. reported that cisplatin-induced NOXA (Phorbol-12-myristate-13-acetate-induced protein 1) is regulated by ATF4 in *TP53*-mutated HNSCC cell lines HN8 (*TP53*-deleted) and HN12 (*TP53*-truncated) (14). As reported in our previous studies, small molecules PG3-Oc and PG3 induce the ISR, ATF4 regulates expression of a subset of proapoptotic p53 target genes, and this leads to apoptosis in *TP53*-mutated or deleted colorectal cancer cells through the HRI-ATF4-PUMA pathway (15, 16). We hypothesized that combination of PG3 plus cisplatin can increase sensitivity to cisplatin in *TP53*-mutated HNSCC cells through enhanced induction of the integrated stress response.

We report that combined treatment of PG3 and cisplatin shows synergistic effects in *TP53*-mutated FaDu (R248L) and Cal27(H193L) HNSCC cells. CReP is a protein that acts as a regulatory subunit of protein phosphatase 1 (PP1) and dephosphorylates eIF2α. Both PG3 and DNA-damage activate HRI, plus degradation of CReP, contribute to the potent induction of eIF2α phosphorylation in PG3 plus cisplatin-treated FaDu and Cal27 cells. Cellular apoptosis induced by the combined PG3 plus cisplatin treatment is regulated through the HRI-ATF4-NOXA pathway. NOXA-mediated Mcl-1 degradation is important for the PG3 plus cisplatin combination treatment-induced apoptosis and is independent of CDK2 kinase in FaDu cells. We observed decreased ATF4 levels after treatments with DNA damage drug cisplatin, CPT, or PG3 plus cisplatin. We verified that cisplatin-induced ATF4 degradation was modulated by kinase CK1δ, CK2, and E3 ligase β-TrCP1. Our results provide a rational strategy of triple treatments, ISR inducer plus DNA damaging drug plus β-TrCP1 inhibitor/CK1δ inhibitor/CK2 inhibitor, to achieve potent and prolonged anti-tumor effects to overcome chemoresistance in HNSCC.

## Results

### Combination of PG3 and cisplatin synergistically inhibits cell viability and enhances the induction of the Integrated Stress Response (ISR)

Cell viability assays were performed to determine whether combined treatment of PG3 and cisplatin is synergistic in HNSCC cells. The cell viability was normalized to the untreated control (100%) as shown in **Figure 1A**. Compared to two normal cell lines, HFF1 (human foreskin fibroblast cell) and IMR90 (human lung fibroblast cell), the combination treatment with PG3 and cisplatin shows more potent inhibitory effects in FaDu and Cal27 HNSCC cells than in HFF1 and IMR90 cells. Also, Cal27 cells are more sensitive to combined PG3 and cisplatin treatment than FaDu cells. The percentage of cell viability, IC50 values, and synergy scores were calculated against the respective DMSO control using the software Combenefit using the model HAS and displayed as surface plots where dose-responses were integrated (17) (**Figure 1B** and **Supplementary Figure 1**).

**Figure 1.**
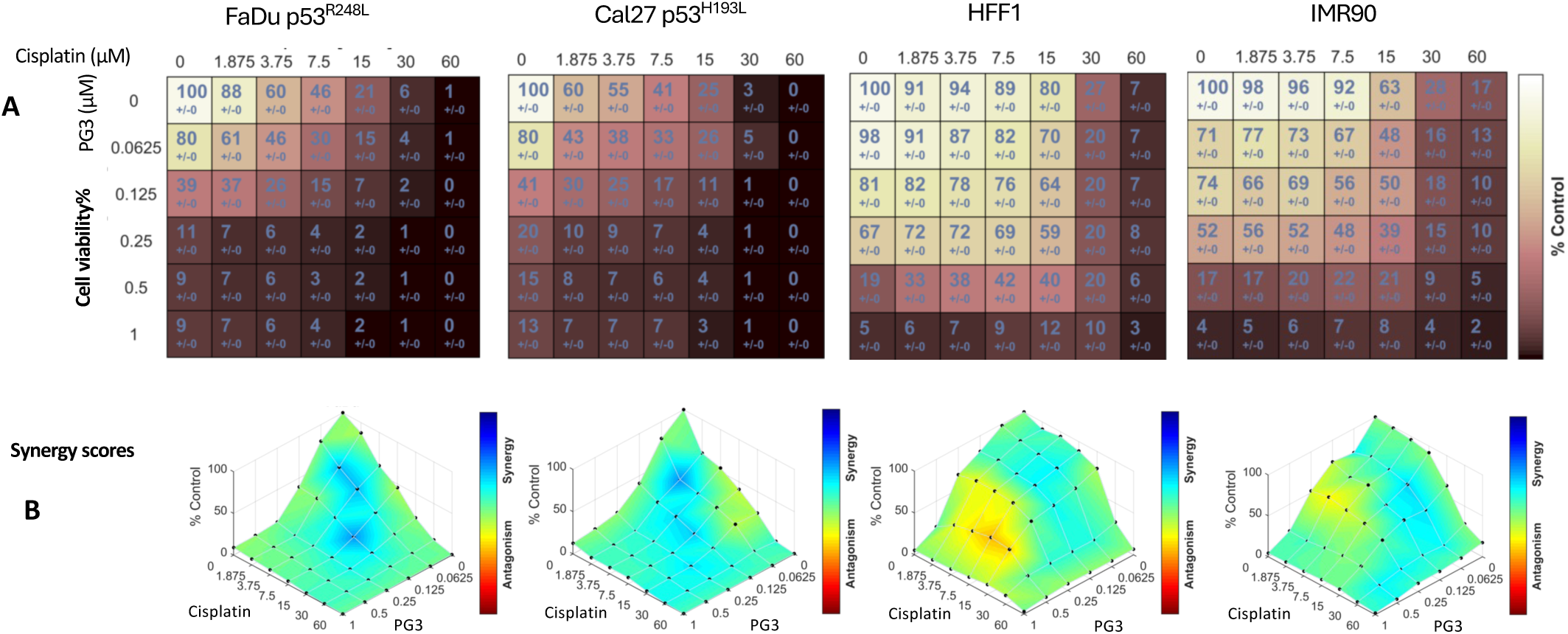
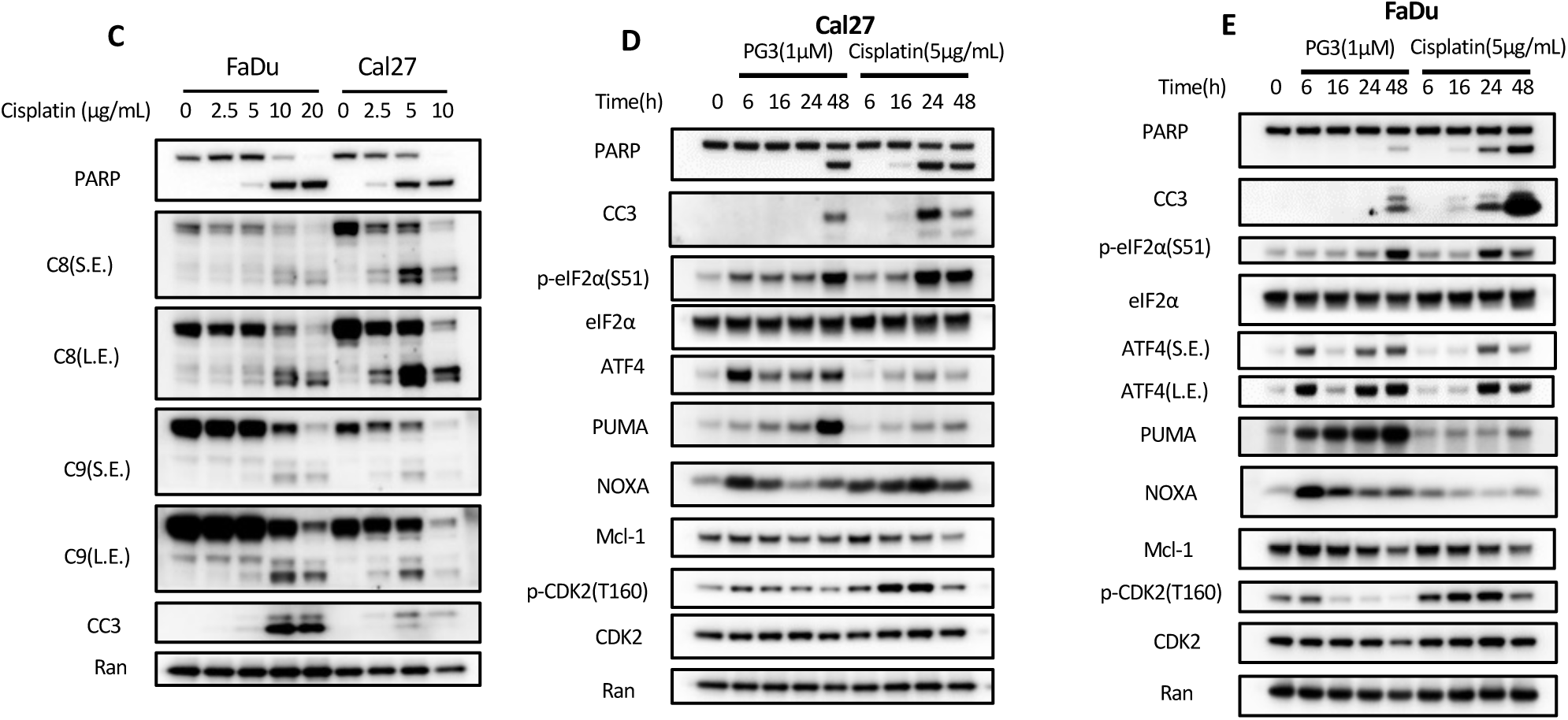
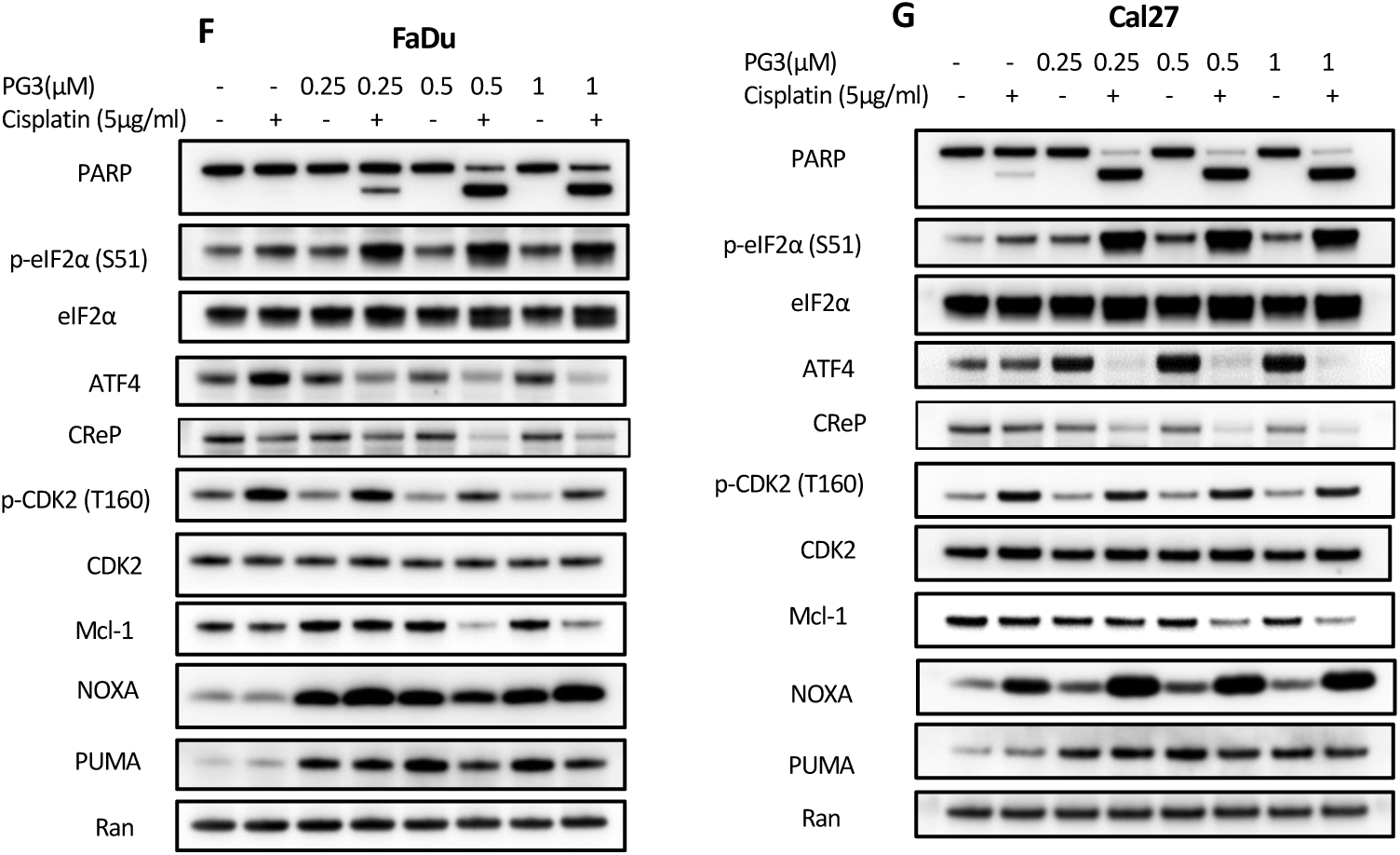
The combination of PG3 and cisplatin synergistically inhibits cell viability and enhances the induction of ISR. **(A)** and **(B)** Cell viability was measured in FaDu, Cal27, HFF-1, and IMR90 cells. Cells were treated with different concentrations of PG3, cisplatin, or DMSO for 72 h. Luciferase activity was imaged using the IVIS Imaging System after treatment. Cell viability data were normalized to those of DMSO treatment control in each cell line, and data analyses were performed using Combenefit software. **(C)** Dose-response experiments: FaDu and Cal27 cells were treated with different doses of cisplatin for 24 hours, then western blot assays were performed using the indicated antibodies. **(D)** and **(E)** Time-course experiments: FaDu and Cal27 cells were treated with PG3 and cisplatin for the indicated periods, respectively, and then western blot assays were performed using the indicated antibodies. **(F)** and **(G)** FaDu and Cal27 cells were treated with PG3, cisplatin, and PG3+cisplatin for 16 hours, and then western blot assays were performed using the indicated antibodies.

At 24-hours of treatment with cisplatin induced apoptosis in a dose-responsive manner in FaDu and Cal27 HNSCC cells as indicated by apoptosis markers, such as cleavage of caspase-8, caspase-9, caspase-3, and PARP (**Figure 1C**). We chose 5 µg/mL (16.7 µM) cisplatin for time-course experiments because at this concentration, cisplatin induced mild caspase-3 and PARP cleavage in Cal27 cells and no caspase-3 cleavage and very minor PARP cleavage in FaDu cells (**Figure 1C**). Time-course experiments were performed with PG3 or cisplatin, respectively. Both PG3 and cisplatin induced phosphorylation of eIF2α on Ser51 in a time-dependent way, indicating the induction of the ISR (**Figure 1D and E**). Both PG3 and cisplatin resulted in upregulation of ATF4, however, PG3 induced a quicker and higher level of ATF4 protein than cisplatin (**Figure 1D and E**). As for ATF4 target genes *PUMA* and *NOXA* expression, PG3 induced strong upregulation of PUMA protein level over time. By contrast, cisplatin showed weak induction of PUMA (**Figure 1D and E**). On the other hand, PG3 induced upregulation of NOXA protein at the 6-hour time point and the protein level then decreased with time. By contrast, cisplatin showed potent induction of NOXA in Cal27 cells but a mild increase at 6-and 48-hour time points in FaDu cells (**Figure 1D and E**).

We chose a 16-hour time point for the combination treatment with PG3 and cisplatin because either PG3 or cisplatin alone induced the cleavages of caspase-3 and PARP were negligible at that time and with the doses used (**Figure 1D and E**). The concentration of cisplatin was maintained at 5 µg/mL while the PG3 concentration was increased from 0.25 µM to 1 µM (**Figure 1F and G**). PG3 or cisplatin alone did not induce PARP cleavage in the FaDu cell (**Figure 1F**). Cisplatin alone, but not PG3, led to very minor cleavage of PARP in the Cal27 cells (**Figure 1G**). However, the combined treatments induced much more cleavage of PARP (**Figure 1F** and **G**). Meanwhile, the combined PG3 and cisplatin treatments induced much higher levels of eIF2α phosphorylation, indicating enhanced induction of the ISR than PG3 or cisplatin alone, which is consistent with our hypothesis.

Although the combined PG3 and cisplatin treatments led to the downregulation of ATF4 protein levels (**Figure 1F** and **G**), the combined treatments induced higher levels of NOXA than with either PG3 or cisplatin alone in the Cal27 cells (**Figure 1G**). In FaDu cells, the combined PG3 and cisplatin treatments maintained slightly higher levels of NOXA than PG3 or cisplatin alone (**Figure 1F**). Our data suggest that ATF4 has completed the upregulation of NOXA before the ATF4 protein level decreased at the 16-hour time point (**Figure 4**). In addition, at the 0.5 and 1 µM concentrations of PG3, the combined PG3 and cisplatin treatments caused potent downregulation of Mcl-1 in both FaDu and Cal27 HNSCC cells (**Figure 1F** and **G**), and more PARP cleavage in FaDu cells than at 0.25 µM PG3 (**Figure 1F**), suggesting the inhibition of Mcl-1 may facilitate cellular apoptosis.

### The ISR mediates cell apoptosis induced by PG3 and cisplatin combined treatment of HNSCC cells

van Galen, *et al*. created an ATF4 mRNA translation reporter (18). We expected that the reporter could be used as an ISR reporter. Bidirectional lentiviral ISR reporter vector was transduced into Cal27 or FaDu HNSCC cells, and the cells were marked by TagBFP (constitutively fluorescent blue fluorescent protein). GFP protein level and brightness measure the induction of the ISR, which is regulated by phospho-eIF2a (**Figure 2A**) (see Materials and Methods for details) (18). Thapsigargin (TG) and ONC201 were known ISR-inducers and used as positive controls to verify the reporter’s activity by fluorescence image (**supplementary Figure 2A**). Although cisplatin or PG3 and cisplatin combination led to ATF4 downregulation, GFP protein levels increased, correlating with increased eIF2α phosphorylation (**Figure 2A** and **supplementary Figure 2B**), indicating a persistence of ISR activation.

**Figure 2.**
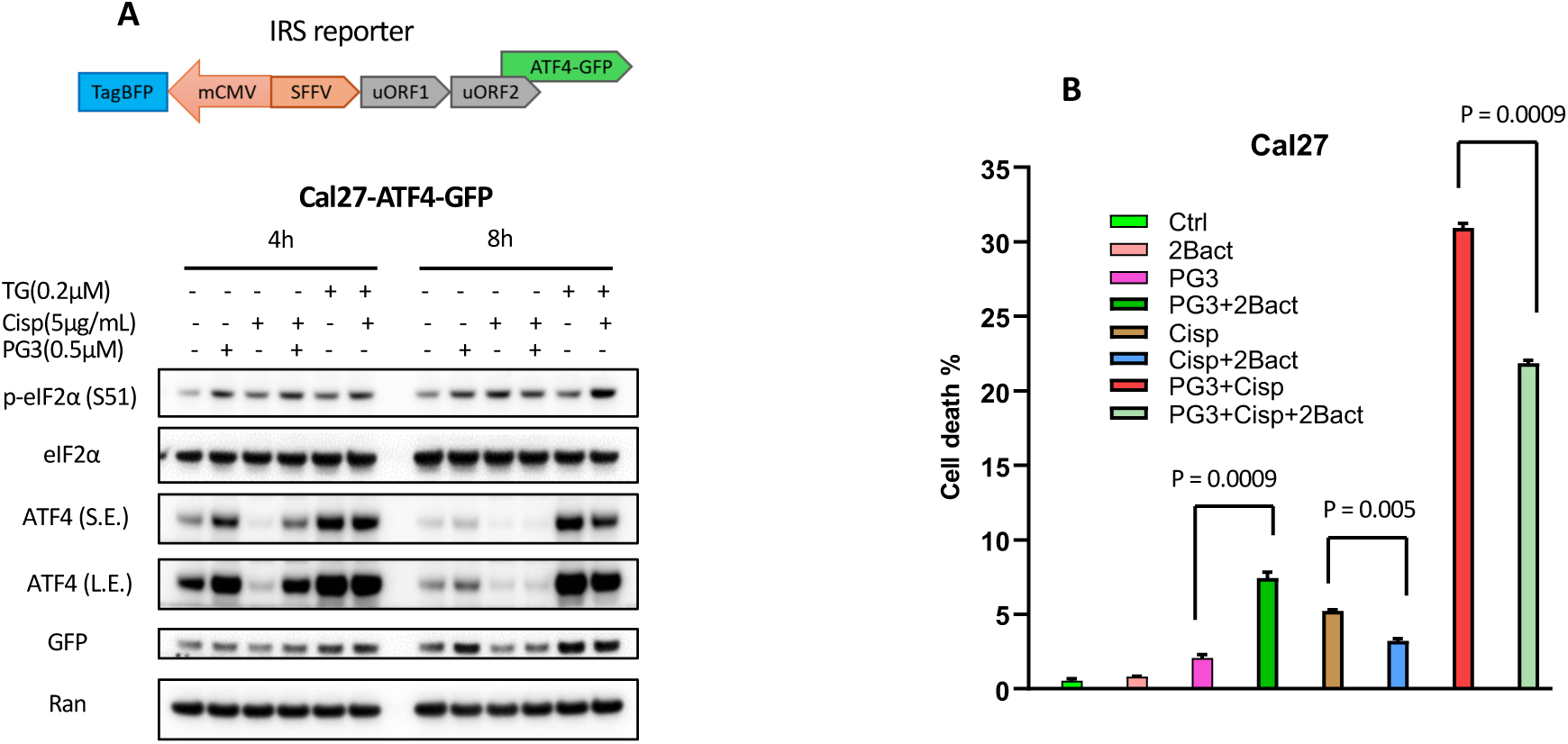
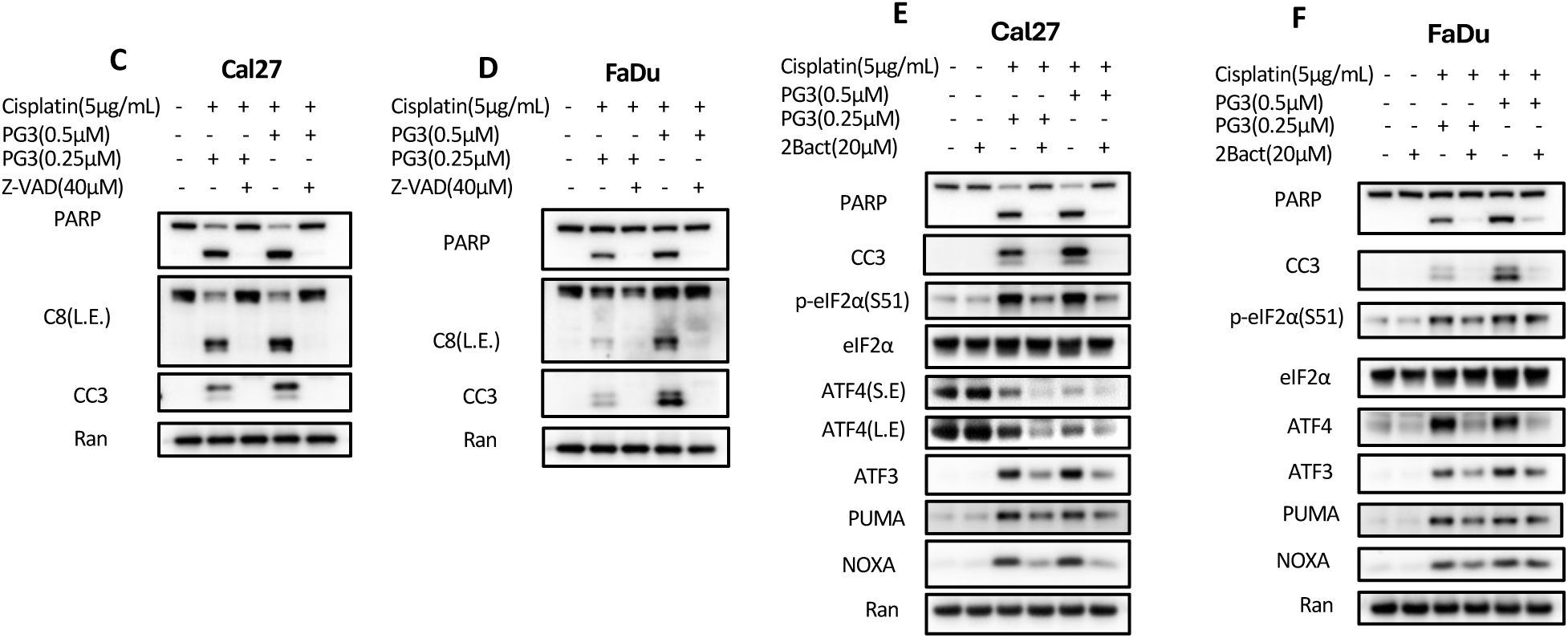
The ISR mediates cell apoptosis induced by the combined treatment. **(A)** Structure of the IRS reporter. The Cal27-ATF4-GFP reporter cells were treated with PG3, cisplatin, and thapsigargin (TG) for 4 or 8 hours, respectively. Western blot was performed using the indicated antibodies. **(B)** Flow cytometric analysis of cell death. Cal27 cells were pre-treated with 2Bact (20 µM) for 30 min, then PG3 (0.5 µM), and cisplatin (5 µg/mL) were added to corresponding wells, and incubated at 37°C for 12 hours. **(C)** and **(D)** Cells were pre-treated with z-VAD-FMK for 30 min, then were treated with the combinations of PG3 plus cisplatin for 16 hours **(C)**, and 21 hours **(D)**, respectively. Western blots were performed using the indicated antibodies. **(E)** and **(F)** Cal27 and FaDu cells were pre-treated with 2Bact (20 µM) for 30 min, then the combined treatments of PG3 plus cisplatin were performed for Cal27 (16 hours), and FaDu (21 hours). Western blot assays were performed using the indicated antibodies.

PG3 is a red fluorescent molecule, and its spectrum overlaps with red fluorescent dyes, such as PI (propidium Iodide), and partially overlaps with green fluorescence dyes. Zombie Violet is a violet fluorescence dye. It distinguishes dead cells by exploiting the permeability of the compromised cell membranes of dead cells, allowing the dye to enter and stain dead cells. 2BAct inhibits the integrated stress response and restores global protein synthesis in cells by activating eIF2B (Eukaryotic translation initiation factor 2B) (19). Wong *et al.* reported that 2BAct prevents neurological defects in mice caused by ISR (19). Flow cytometric analysis showed that 2Bact partially inhibited the cell death induced by PG3 and cisplatin (**Figure 2B** and **supplementary Figure 2C**) as the dye stains late-stage death of cells and does not identify early apoptotic cells, suggesting the PG3 and cisplatin also causes other types of cell death.

The combination treatment with various concentrations of PG3 showed potent PARP cleavage, suggesting apoptosis was induced by the PG3 and cisplatin combination treatments compared to PG3 or cisplatin alone (**Figure 1F** and **G**). To verify this observation, a pan-caspase inhibitor (z-VAD-FMK) was used. z-VAD-FMK blocked PG3 and cisplatin combination treatment-induced cleavage of caspase-8, caspase-3, and PARP, consistent with apoptosis induction in FaDu and Cal27 HNSCC cells (**Figure 2C** and **D**). As shown in **Figure 2E** and **F**, 2Bact reduced the PG3 and cisplatin combined treatment-induced eIF2α phosphorylation and potently inhibited upregulation of ATF4 and its target gene expression of ATF3, PUMA, and NOXA. Moreover, 2Bact abolished the PG3 and cisplatin combination treatment-induced cleavage of caspase-3 and PARP, indicating that the ISR mediates the PG3 and cisplatin combined treatment-induced cellular apoptosis.

### NOXA is required for the combined PG3 and cisplatin treatment-induced apoptosis of HNSCC cells

NOXA and PUMA are proapoptotic BH3-only proteins and known ATF4 target genes (11, 15, 16, 20). Sharma, *et al.* reported that cisplatin led to NOXA-mediated apoptosis in p53-deficient HNSCC HN8 and HN12 cells (14). We investigated whether the cisplatin-induced NOXA mediates apoptosis in *TP53*-mutated FaDu and Cal27 HNSCC cells. siRNA knockdown of NOXA was efficient (**Figure 3A** and **B**) and potently suppressed cleavage of caspase-8, caspase-9, caspase-3, and PARP in FaDu and Cal27 HNSCC cells, indicating that NOXA plays an important role in cellular apoptosis under the experimental conditions (**Figure 3A** and **B**). Surprisingly, silencing PUMA led to increased cleavage of PARP, caspase-8 and caspase-3 (**Figure 3A** and **B**), suggesting that PUMA antagonized cisplatin-induced apoptosis.

**Figure 3.**
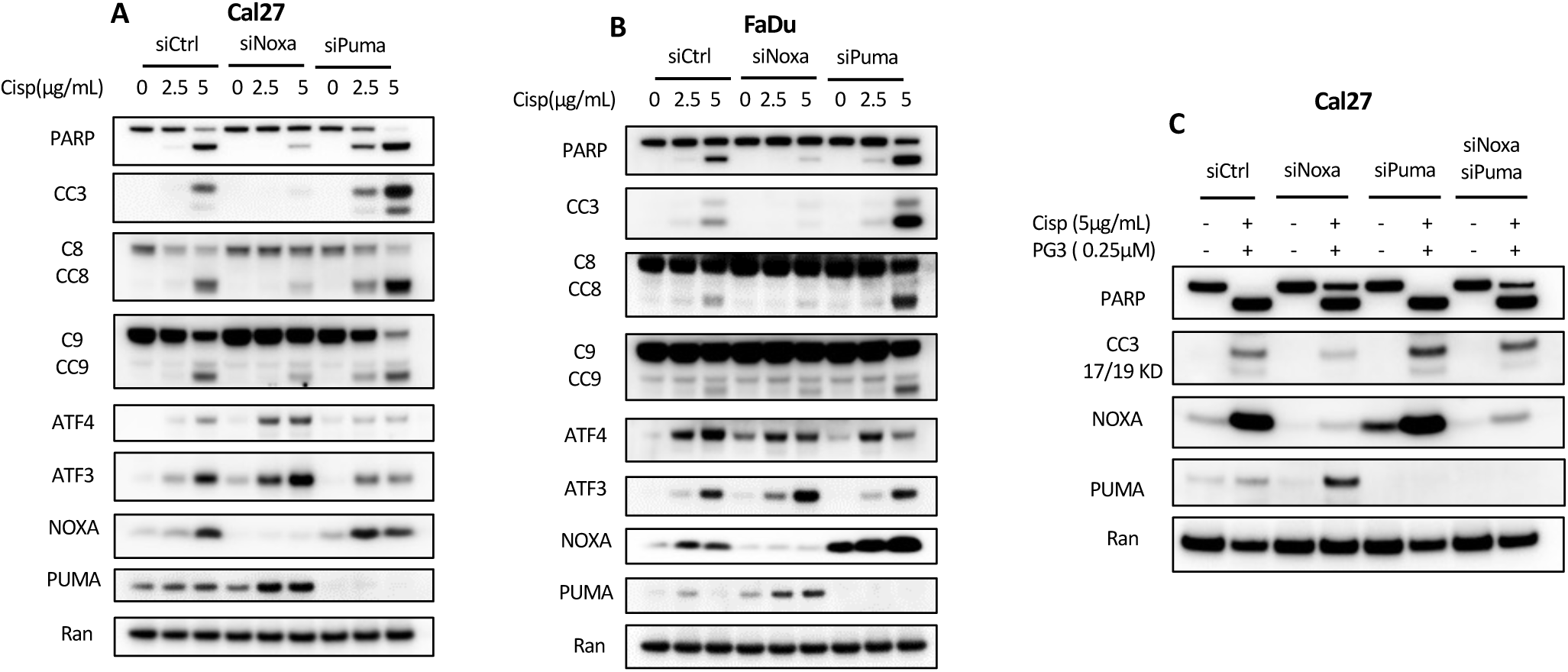
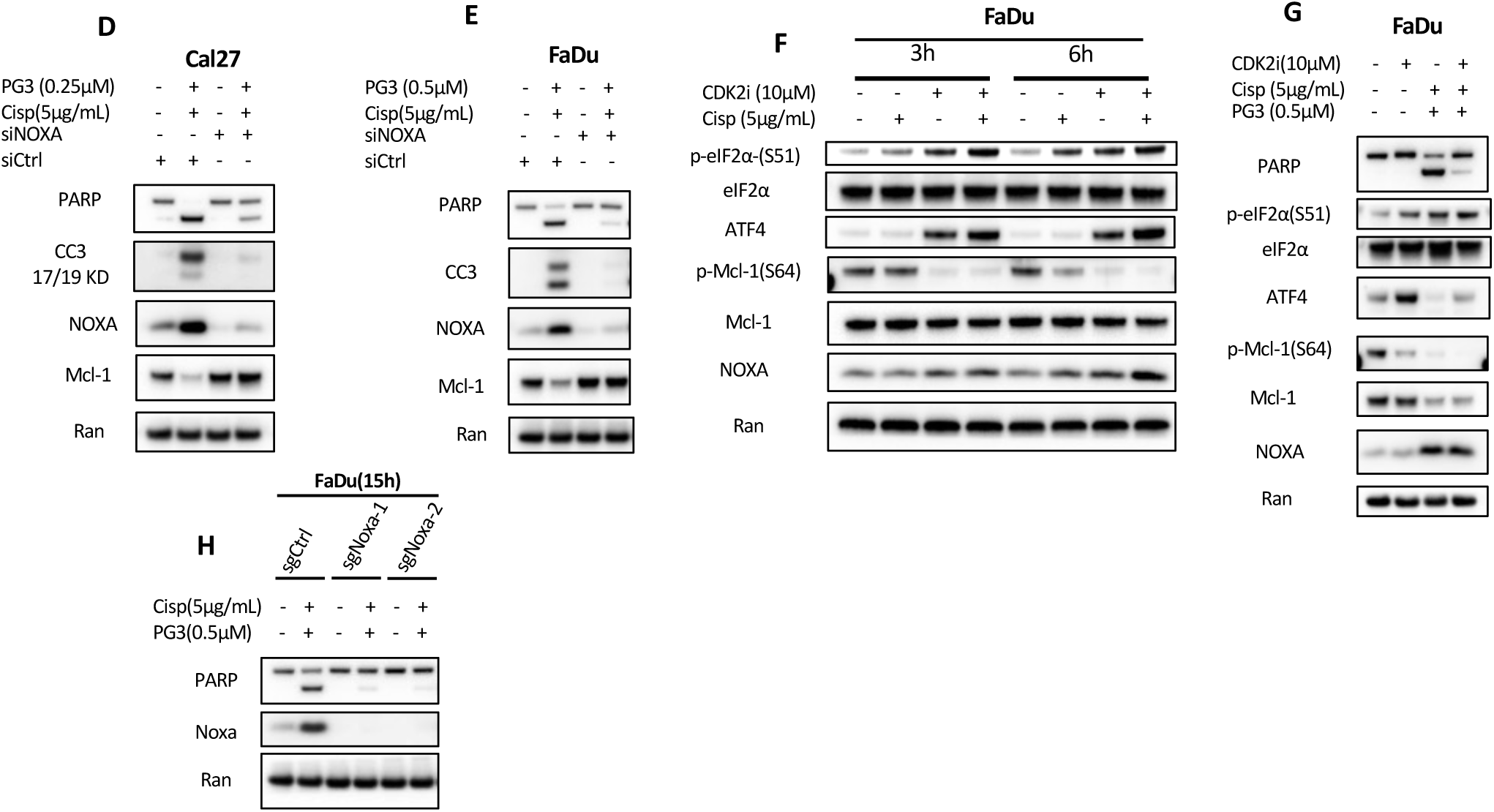
NOXA-mediated Mcl-1 degradation is independent of CDK2. **(A)** Ca27 and **(B)** FaDu HNSCC cells were transfected with Control, NOXA, or PUMA siRNAs. At 24 hours after transfection, the Cal27 and FaDu cells were treated with cisplatin for 15 or 22 hours, respectively. Then, cell lysates were prepared, and western blot analysis was performed using the indicated antibodies. **(C)** Cal27 cells were transfected with Control, NOXA, or PUMA siRNAs. After 24 hours, the cells were treated with PG3 plus cisplatin for 17 hours. Then, western blot analysis was performed using the indicated antibodies. **(D)** and **(E)** Cal27 and FaDu cells were transfected with Control, NOXA siRNAs for 24 hours, and then the cells were treated with the combinations of PG3 plus cisplatin for 12 hours. Western blot analysis was performed using the indicated antibodies. **(F)** FaDu cells were pre-treated with the CDK2 inhibitor (CDK2i) for 1 hour, and then the cells were treated with cisplatin for 3 or 4 hours, respectively. Western blot assay was performed using the indicated antibodies. **(G)** FaDu cells were pre-treated with the CDK2i for 1 hour, then treated with PG3 plus cisplatin for 15 hours. Western blot assay was performed using the indicated antibodies.

PG3-induced PUMA-mediated apoptosis in *TP53*-mutated colorectal cancer cell lines HT29 (R273H) and HCT116 (*TP53^-/-^*) (16). The combined PG3 and cisplatin treatment induced mild upregulation of PUMA and potent upregulation of NOXA (**Figure 1F** and **G**). We investigated whether the PG3 and cisplatin combined treatment leads to apoptosis through NOXA, PUMA, or both NOXA and PUMA in HNSCC cells. As shown in **Figure 3C**, the combined PG3 and cisplatin treatment caused strong cleavage of PARP, and full-length PARP was not detected in control-siRNA-treated Cal27 cells, suggesting potent apoptosis. siRNA knockdown of NOXA reduced caspase-3 cleavage and partially restored the protein level of full-length PARP compared to control siRNA, indicating partial protection of Cal27 cells from NOXA-induced apoptosis. Silencing PUMA increased cleavage of caspase-3 compared to control siRNA and did not lead to any level of restoration of full-length PARP, indicating that its silencing promoted cellular apoptosis. Double knockdown of both NOXA and PUMA restored the protein level of full-length PARP to almost the same level as the knockdown of NOXA alone, suggesting that NOXA mediated the combination treatment-induced apoptosis (**Figure 3C**).

Unexpectedly, the knockdown of PUMA increased NOXA-induced apoptosis (**Figure 3A, B**, and **C**). These observations are consistent with publications summarized in the review article (21). PUMA competes with NOXA binding to Mcl-1 (Myeloid cell leukemia-1) (21). Knockdown of PUMA would facilitate NOXA binding to Mcl-1. Second, because binding of PUMA to Mcl-1 blocks E3-ligase Mule binding to Mcl-1 (21), knockdown of PUMA should facilitate NOXA-mediated Mule binding to Mcl-1 and Mule-mediated ubiquitination and degradation of Mcl-1.

### NOXA-mediated Mcl-1 degradation is independent of CDK2 in PG3 and cisplatin-treated HNSCC cells

Mcl-1 is an anti-apoptotic protein and a member of the Bcl-2 family, playing a crucial role in preventing cell death, and is often overexpressed in cancers (22). NOXA binds and mediates Mcl-1 degradation (22). Degradation of Mcl-1 leads to mitochondrial outer membrane permeabilization (MOMP) and the subsequent release of factors like cytochrome c, which initiates the apoptotic cascade *via* cleavage and activation of caspase-3 and PARP (23). Since the combined PG3 and cisplatin treatments led to potent upregulation of NOXA and downregulation of Mcl-1 (**Figure 1F** and **G**), we investigated whether NOXA mediated Mcl-1 degradation. As shown in **Figure 3D** and **E**, the knockdown of NOXA abolished the co-treatment-induced degradation of Mcl-1, and PARP cleavage, indicating that NOXA mediates PARP cleavage and Mcl-1 degradation. Meanwhile, restoration of Mcl-1 protein level correlated well with rescued apoptosis as indicated by blocking the cleavage of caspase-3 and PARP in both FaDu and Cal27 HNSCC cells (**Figure 3D** and **E**). Knocking out the *NOXA* gene potently blocked PG3 and cisplatin co-treatment-induced PARP cleavage (**Figure 3H**), further confirming that NOXA mediated the co-treatment-induced apoptosis.

CDK2 (Cyclin-dependent kinase 2) is a key kinase that is involved in the regulation of cell cycle progression. It plays a key role in the G1/S phase transition (24, 25). Bim (Bcl-2 interacting mediator of cell death) is a pro-apoptotic BH3-only protein (24, 25). CDK2-dependent phosphorylation of Mcl-1 residue Ser64 increases Bim binding and enhances Mcl-1 anti-apoptotic function (24–26). In addition, Choudhary *et al.* reported that CDK2 phosphorylates Mcl-1 residues (Thr92 and Thr163) and stabilizes Mcl-1 (24). On the contrary, it has been shown that cisplatin-induced Mcl-1 degradation is mediated by both NOXA and CDK2-regulated Mcl-1 phosphorylation (Ser64/Thr70) in HeLa cells (27).

To investigate whether cisplatin activates CDK2 in FaDu and Cal27 HNSCC cells, a western blot was performed using phospho-CDK2(T160) specific antibody; phosphorylation of CDK2 residue Thr160 was increased with time, indicating CDK2 activation (**Figure 1D** and **E**). The combination treatment of PG3 and cisplatin also activated CDK2 as potently as cisplatin (**Figure 1F** and **G**). A time-course experiment indicated that cisplatin alone did not result in downregulation of Mcl-1 within 24 hours (**Figure 1D** and **E**). Cisplatin or PG3 and cisplatin inhibited phosphorylation of Mcl-1 (Ser64) (**Figure 3F** and **G**), though CDK2 was activated, suggesting that CDK2 did not phosphorylate Mcl-1 on Ser64. CDK2 inhibitor II (CDK2i) is a CDK2-selective inhibitor (27). CDK2i suppressed phosphorylation of Mcl-1 Ser64, however, CDK2i did not rescue degradation of Mcl-1 protein induced by PG3 and cisplatin (**Figure G**), which is consistent with the results reported by others; phosphorylation of Mcl-1 residue Ser64 does not regulate Mcl-1 protein stability and instead enhances Mcl-1 anti-apoptotic function (24–26).

Collectively, although cisplatin activates CDK2 in FaDu and Cal27 HNSCC cells, cisplatin leads to dephosphorylation of Mcl-1 (S64) instead of phosphorylation. Consistent with this observation, cisplatin-caused NOXA-mediated Mcl-1 degradation is independent of CDK2 in the FaDu and Cal27 cells.

### ATF4 is required for upregulation of NOXA and apoptosis in PG3 and cisplatin-treated HNSCC cells

Sharma, *et al*. reported that ATF4 and ATF3 regulated cisplatin-induced NOXA in *TP53*-deficient HN8 and HN12 cells (14). Since the combination of PG3 and cisplatin induced potent ISR activation in *TP53*-mutated FaDu and Cal27 HNSCC cells, we investigated whether ATF4 and ATF3 modulate the upregulation of NOXA. At the 6-hour timepoint, siRNA knockdown of ATF3 did not inhibit NOXA upregulation induced by PG3, cisplatin, or PG3 and cisplatin, respectively. However, knockdown of ATF4 suppressed both NOXA and ATF3 upregulation induced by PG3, cisplatin, and PG3 and cisplatin, respectively, indicating that ATF4 regulates NOXA and ATF3 expression (**Figure 4A** and **B**). At the 12-hour treatment, the silencing of ATF3 did not decrease NOXA upregulation induced by cisplatin or PG3 and cisplatin and did not lead to restoration of Mcl-1 protein and reduction of PARP cleavage (**Figure 4C** and **D**). By contrast, silencing ATF4 reduced NOXA induction in Cal27 cells but not in FaDu HNSCC cells. Importantly, the knockdown of ATF4 led to the restoration of Mcl-1 protein levels and reduction of PARP cleavage in both Cal27 and FaDu HNSCC cells, suggesting that the ATF4-NOXA-Mcl-1 pathway modulates apoptosis (**Figure 4C** and **D**). Compared to the 6-hour timepoint (**Figure 4A** and **B**), after 12-hour treatment, ATF3 and NOXA protein levels were restored through unknown mechanisms, even if silencing ATF4 was efficient (**Figure 4C** and **D**). Overall, ATF4 is required for upregulation of NOXA, and the ATF4-NOXA-Mcl-1 axis regulates apoptosis induced by PG3 and cisplatin treatment.

**Figure 4.**
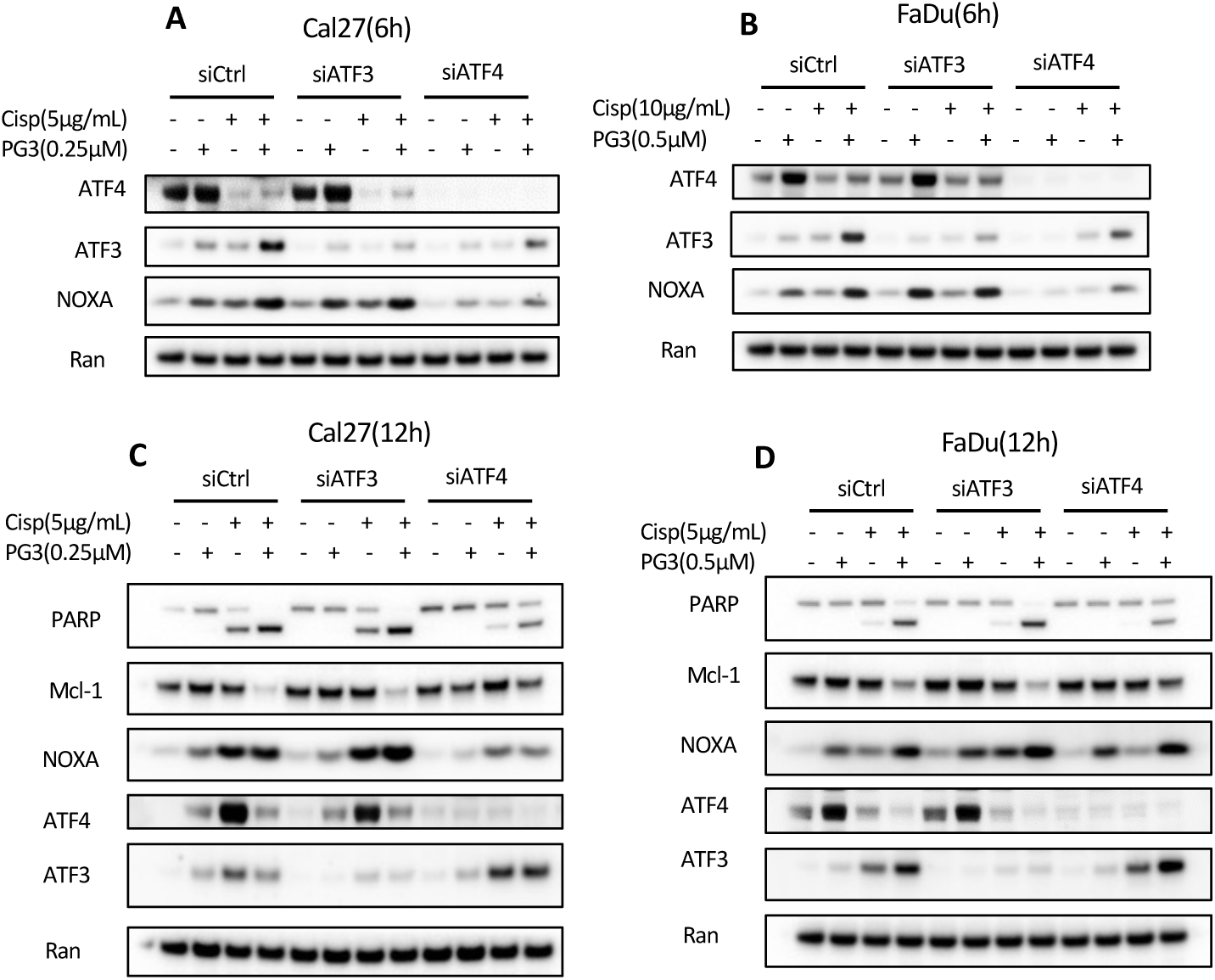
ATF4 is required for upregulation of NOXA and cell apoptosis. **(A)** Cal27 and **(B)** FaDu cells were transfected with Control, ATF3, and ATF4 siRNAs, respectively. At 24 hours after transfection, the Cal27 and FaDu cells were treated with cisplatin for 6 hours, respectively. Then, cell lysates were prepared, and western blot analysis was performed using the indicated antibodies. **(C)** Cal27 and FaDu **(D)** cells were transfected with Control, ATF3, or ATF4 siRNAs. After 24 hours of the transfection, the Cal27 or FaDu cells were treated with cisplatin for 12 hours, respectively. Western blot analysis was performed using the indicated antibodies.

### Cisplatin activates kinase HRI in cancer cells

Differential DNA damage activation of eIF2α kinases is dependent on both DNA damaging agents and cell type (12, 13, 28–33). UV radiation leads to DNA damage-mediated GCN2 activation (12, 13, 28–30). XRCC1 (X-ray repair cross-complementing protein 1) is crucial for DNA single-strand break repair (SSBR) and base excision repair (BER). Single-strand breaks (SSBs) caused by the downregulation of scaffold protein XRCC1 activate PERK (32). Mitochondrial DNA double-strand breaks (mtDSBs) activate HRI (33).

To determine which eIF2α kinase is activated by cisplatin, *PERK, GCN2*, *PKR*, or *HRI*-knockout cell lines were treated with cisplatin for 16 hours (**Figure 5A**). PANC1 *HRI^-/-^* and PANC1 *PKR^-/-^* cell lines were created using number 2 sgHRI-2 and number 1 sgPKR-1 respectively and were verified in our published article (16). Two *PERK*-KO and two *GCN2*-KO PANC1 cell lines were used. Only knockout of the *HRI* gene blocked cisplatin-induced eIF2α phosphorylation (**Figure 5A**). To confirm this observation, PANC1 *HRI^-/-^* cells were treated with cisplatin for 6 hours, western blots showed that knockout of *HRI* gene blocked cisplatin (5 µg/ml) induced eIF2α phosphorylation (**Figure 5B**), consistent with the 16-hour cisplatin (5µg/ml) treatment result (**Figure 5A**).

**Figure 5.**
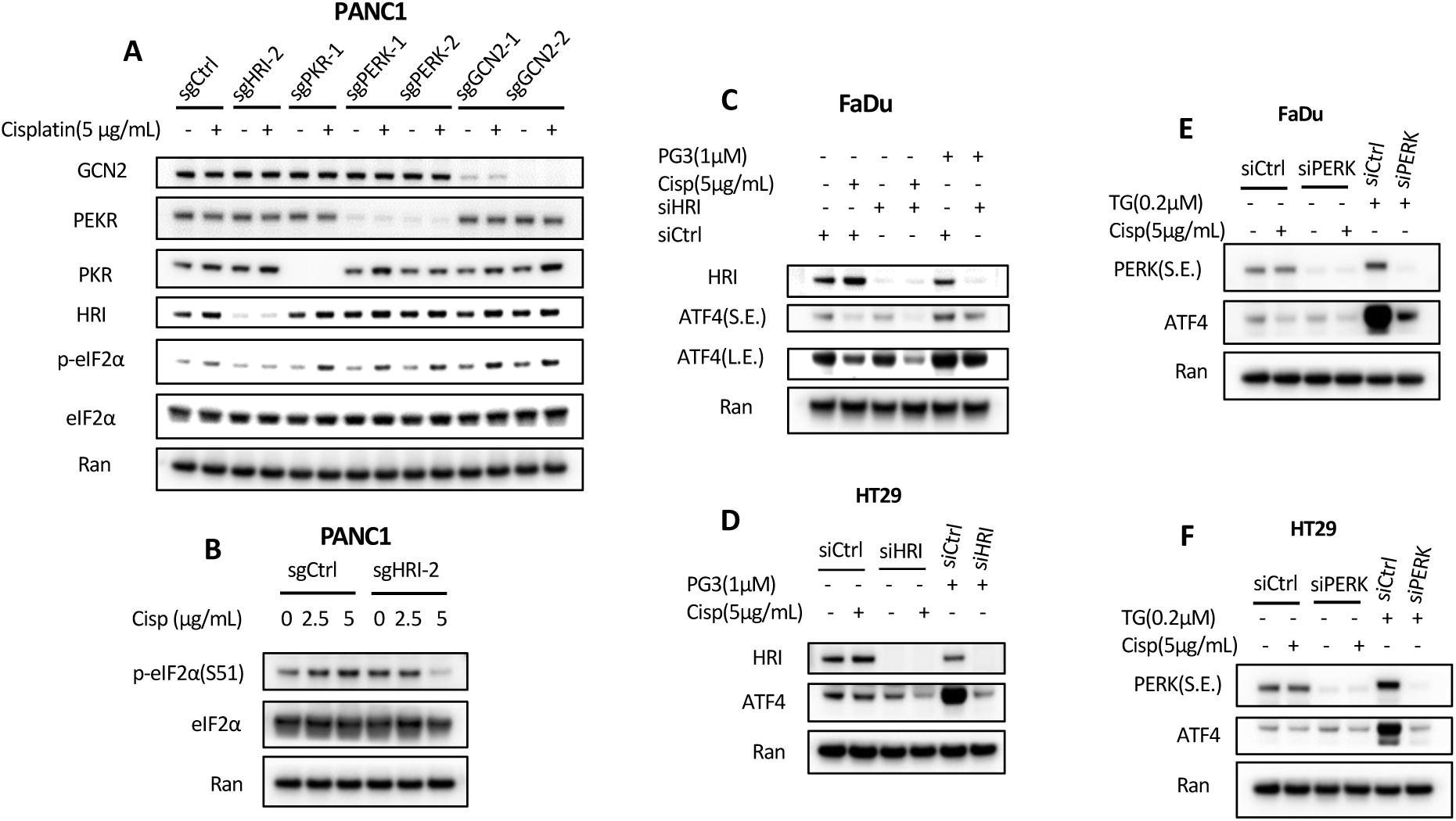
Cisplatin activates HRI. **(A)** sgControl PANC1, PANC1 *HRI^-/-^* #2, PANC1 *PKR^-/-^* #1, PANC1 *PERKI^-/-^* #1/2, and PANC1 *GCN2^-/-^* #1/2 cells were treated with cisplatin for 16 hours, then a western blot assay was performed using the indicated antibodies. **(B)** sgControl PANC1, and PANC1 *HRI^-/-^* #2 cells were treated with cisplatin for 6 hours, then a western blot assay was performed using the indicated antibodies. **(C)** FaDu and **(D)** HT29 cells were transfected with Control or HRI siRNAs. After 24 hours, the FaDu and HT29 cells were treated with cisplatin or PG3 for 16 hours. Western blot analysis was performed using the indicated antibodies. **(E)** FaDu and **(F)** HT29 cells were transfected with Control or PERK siRNAs. After 24 hours’ transfection, the FaDu and HT29 cells were treated with cisplatin and TG for 15 or 21 hours, respectively. Western blot analysis was performed using the indicated antibodies.

To determine whether cisplatin activates HRI in FaDu and HT29 cells, siRNA knockdown of *HRI* was performed using the siHRI RNA that was verified in our previous publication (16). PG3 was included as a positive control because PG3 activates HRI (16). Knockdown of *HRI* suppressed cisplatin-or PG3-induced ATF4 compared to control siRNA (**Figure 5C** and **D**), suggesting cisplatin activates HRI in both FaDu and HT29 cells.

We observed that both single guide RNA sgPERK-1 and sgPERK-2 did not achieve a highly effective knockout of *PERK,* as the western blot detected PERK signal (**Figure 5A**). To determine whether PERK could also be activated by cisplatin, silencing of *PERK* was performed, and thapsigargin (TG) was included as a positive control for PERK activation because TG induces ER stress and activates PERK (34). Knockdown of *PERK* did not show any effects on ATF4 protein levels treated with cisplatin, while blocking TG-induced ATF4 upregulation (**Figure 5E** and **F**). TG treatment led to PERK band upshift on the gels compared to untreated controls, indicating PERK activation in FaDu cells (**Figure 5E**) and HT29 cells (**Figure 5F**). By contrast, cisplatin didn’t cause a PERK band upshift as compared to untreated controls, indicating cisplatin did not lead to PERK activation (**Figure 5E** and **F**). Taken together, gene knockout and siRNA silencing reveal that cisplatin activates HRI in FaDu, HT29, and PANC1 cells.

### Degradation of both ATF4 and CReP is mediated by E3 ligase β-TrCP1

β-TrCP1 (β-transducin repeat-containing protein 1) is an F-box protein that serves as a substrate recognition subunit for the SCF (Skp1-Cullin1-F-box) β-TrCP1 E3 ubiquitin ligase complex (35). ATF4 (36, 37) and CReP (30) are known β-TrCP1 substrates. UV and camptothecin (CPT) cause DNA damage and β-TrCP1-mediated degradation of CReP, resulting in the accumulation of phospho-eIF2α and ISR (29, 30). ATF4 is an unstable protein with a half-life of about 30 minutes under normal conditions (37). Phosphorylation of ATF4 (S219) mediated by kinase CK1δ (Casein kinase 1δ) (6, 36) is essential and primes ATF4 phosphorylation at S215 by kinase GSK3β (38) or CK2 (Casein Kinase 2) (39). Together, the phosphorylation of ATF4 at both S215 and S219 creates a docking site for β-TrCP1 binding and β-TrCP1-mediated ubiquitination and degradation.

Because PG3 induced upregulation of ATF4 but did not lead to downregulation of ATF4 later (**Figure 1F** and **G**), we investigated whether cisplatin leads to ATF4 degradation in the combination treatments of PG3 and cisplatin (**Figure 1F** and **G**). In addition, we observed that the combination of PG3 and cisplatin induced a much higher level of eIF2α phosphorylation than PG3 or cisplatin alone (**Figure 1F** and **G**). Since cisplatin also causes DNA damage, we asked whether cisplatin could lead to CReP degradation and contribute to enhanced eIF2α phosphorylation.

Pevonedistat (MLN4924) is a first-in-class NEDD8-activating enzyme (NAE) inhibitor. Conjugation of NEDD8 to the scaffold cullin1 protein in the SCF complex is carried out by NAE, and is required to activate β-TrCP1 (40). Pevonedistat induced the accumulation of ATF4 protein and led to upregulation of NOXA without increasing eIF2α phosphorylation compared to the untreated controls, indicating that pevonedistat leads to the upregulation of ATF4 through inhibition of β-TrCP, and is independent of the ISR (**Figure 6A** and **B**). FaDu and Cal27 cells were treated for 4 and 7 hours (**Figure 6A** and **B**), and CPT was included as a positive control for DNA-damage-induced degradation of CReP. Pevonedistat rescued cisplatin-induced degradation of ATF4 and led to increased induction of NOXA than cisplatin alone (**Figure 6A** and **B**). CPT did not lead to upregulation of NOXA (**Figure 6A** and **B**). Pevonedistat also rescued CPT-induced degradation of ATF4 but did not lead to the induction of NOXA. In particular, the stabilization of CReP by pevonedistat reduced cisplatin-or CPT-induced phospho-eIF2α, suggesting that degradation of CReP contributed to cisplatin-induced accumulation of eIF2α phosphorylation (**Figure 6A** and **B**). As shown before, the combination of PG3 and cisplatin resulted in potent downregulation of CReP, which is well correlated to the PG3 and cisplatin combination treatment-induced higher level of eIF2α phosphorylation versus PG3 or cisplatin alone (**Figure 1F** and **G**). Collectively, these analyses suggest that β-TrCP1 mediates cisplatin-induced degradation of ATF4 and CReP, and the latter, together with the activation of HRI by both PG3 and cisplatin, contributes to cisplatin-induced potent eIF2α phosphorylation. In summary, A model for the PG3 and cisplatin combination-induced potent ISR and related HRI-ATF4-NOXA signaling pathway is proposed (**Figure 6C**).

**Figure 6.**
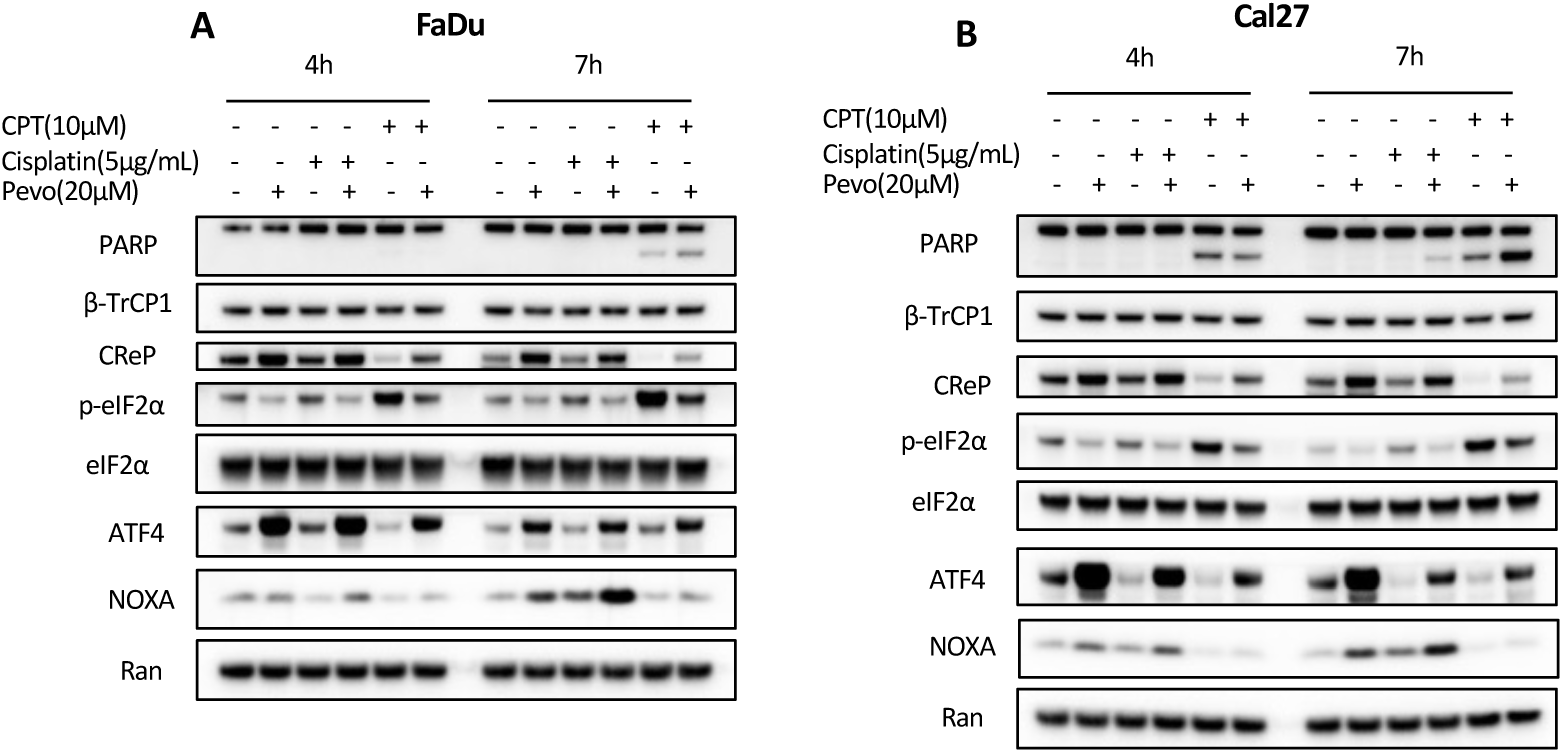
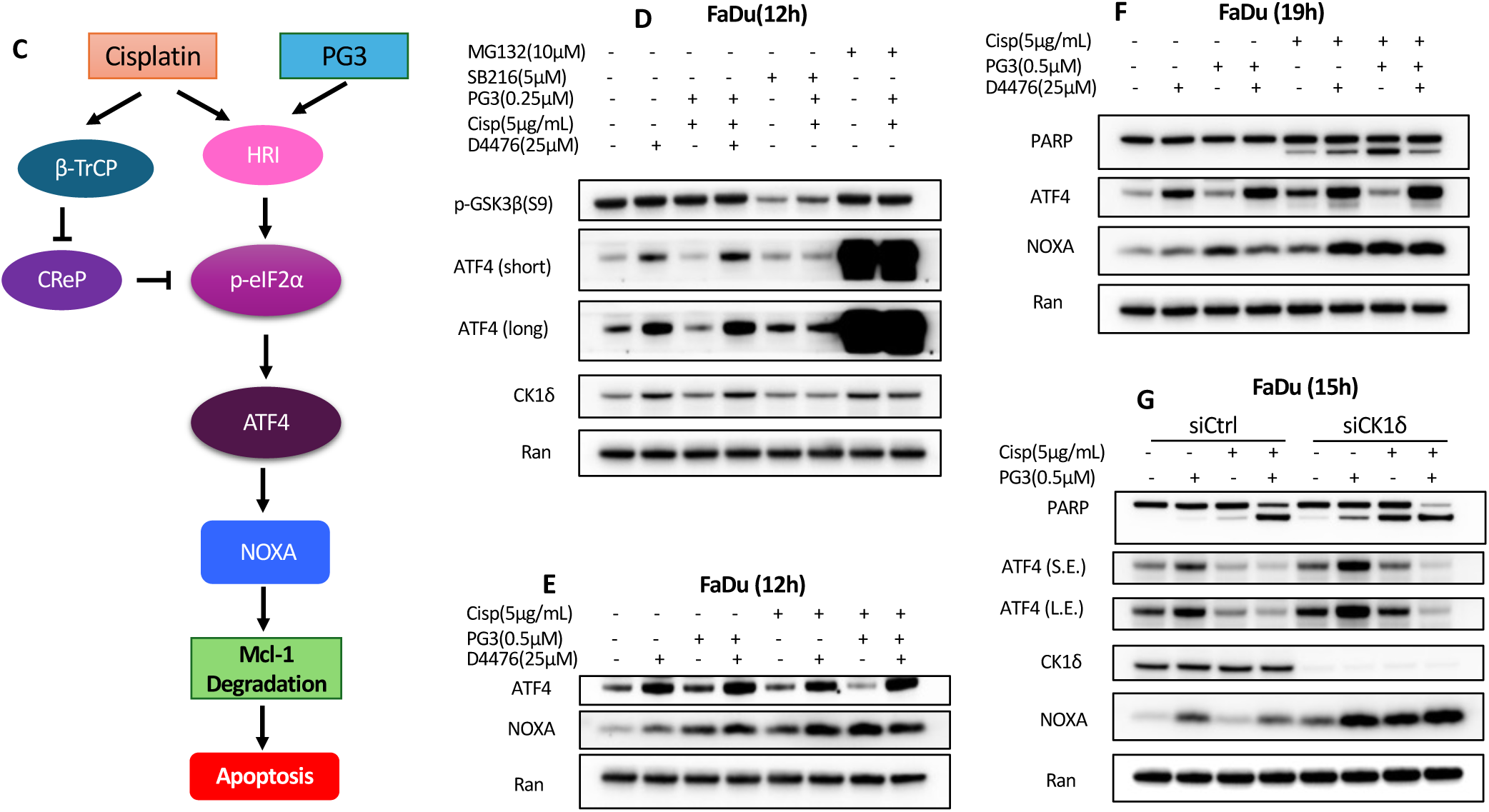
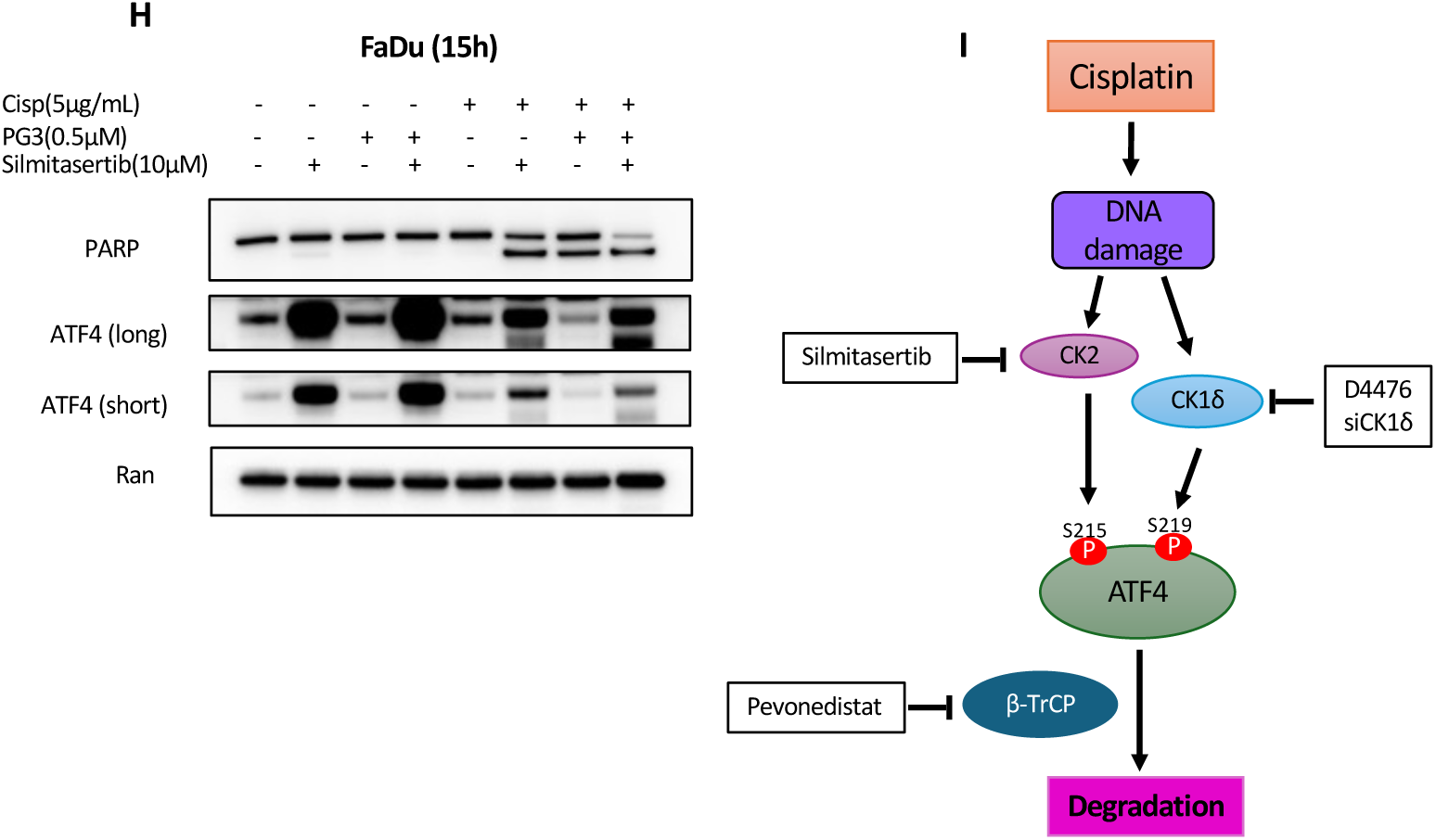
Degradation of ATF4 and CReP is mediated by E3 ligase β-TrCP1 via kinase CK1δ and CK2. **(A)** and **(B)** cells were pre-treated with pevonedistat for 1 hour, and then cisplatin, or CPT was added to the solution for 4 or 7 hours, respectively. Western blot analysis was performed using the indicated antibodies. **(C)** Proposed signaling pathway that regulates the apoptosis induced by the PG3+cisplatin treatment. **(D)** FaDu cells were pre-treated with D4476, SB216763, and MG132 for 1 hour, then continued to be treated with PG3, cisplatin, and PG3 plus cisplatin, respectively, for 12 hours. Western blot analysis was performed using the indicated antibodies. **(E)** And **(F)** FaDu cells were pre-treated with D4476 for 1 hour, then treated with PG3, cisplatin, and PG3 plus cisplatin, respectively, for 12 hours **(E)**, and 19 hours **(F)**. Western blot analysis was performed using the indicated antibodies. **(G)** FaDu cells were transfected with Control and CK1δ siRNAs, respectively. After 24 hours of transfection, the cells were treated with PG3, cisplatin, or PG3 plus cisplatin, for 15 hours. Western blot assay was performed using the indicated antibodies. **(H)** FaDu cells were pre-treated with silmitasertib for 1 hour, and then treated with PG3, cisplatin, or PG3 plus cisplatin, for 15 hours. Western blot analysis was performed using the indicated antibodies. **(I)** Proposed model for cisplatin-induced degradation of ATF4 protein.

### DNA damage-induced degradation of ATF4 is regulated by kinases CK1δ and CK2

DNA damage activates kinase CK1δ (41). Activated CK1δ not only phosphorylates p53 to stabilize it, but also phosphorylates MDM2, leading to its degradation (41). D4476 is a CK1δ inhibitor, and SB216763 is a GSK3α/β inhibitor. D4474 alone stabilized ATF4 compared to the untreated control (**Figure 6D**). Treatment of PG3 and cisplatin led to degradation of ATF4, which could be rescued by D4476 (**Figure 6D**). But SB216763 did not show any stabilization effect on ATF4, and it did not rescue the PG3 and cisplatin combination treatment-induced degradation of ATF4 (**Figure 6D**), indicating that GSK3β does not regulate cisplatin-induced ATF4 degradation in FaDu cells. Proteasome inhibitor MG132 potently stabilized ATF4, which is consistent with ATF4 degradation through the proteasome (**Figure 6D**).

To investigate whether ATF4 stabilized by D4476 could induce more NOXA, western blots were performed in FaDu cells for different periods (**Figure 6E** and **F**). For 12-hours of treatment, co-treatment of cisplatin and D4467 induced more ATF4 and NOXA proteins than cisplatin alone. Although triple-treatment of PG3 plus cisplatin and D4467 led to higher levels of ATF4 than PG3 and cisplatin, the level of NOXA was almost the same (**Figure 6E**). For the 19-hour treatment, the co-treatment of cisplatin and D4467 continued to induce more ATF4 and NOXA protein and led to more PARP cleavage than cisplatin (**Figure 6F**). Triple-treatment of PG3 plus cisplatin and D4467 led to much higher levels of ATF4 than PG3 and cisplatin, however, the level of NOXA protein was the same (**Figure 6E**). Surprisingly, the triple-treatment showed some protection of cells from apoptosis as indicated by weaker PARP cleavage than with PG3 and cisplatin (**Figure 6F**), which is possibly due to D4476 off-target effects. To verify this, siRNA silencing of CK1δ was performed (**Figure 6G**). CK1δ silencing stabilized ATF4, increased the induction of NOXA and PARP cleavage in cells treated with PG3 or cisplatin compared to control siRNA, indicating more cellular apoptosis. As for the combination treatment of PG3 and cisplatin, knockdown of CK1δ induced potent upregulation of NOXA than control siRNA. The cellular apoptosis after CK1δ silencing was much more pronounced, as indicated by a very weak full-length PARP signal, though the levels of cleaved PARP were almost the same as control siRNA, suggesting the triple treatment of PG3 plus cisplatin and CK1δ inhibitor is more potent due to stabilization of ATF4 (**Figure 6G**). During our work, we observed that when cells are experiencing a strong apoptosis, usually the ATF4 protein is hardly detectable (**Figure 6G**). In summary, both siRNA silencing and pharmacological inhibition of CK1δ demonstrated that CK1δ regulates degradation of ATF4, which is consistent with the work of others (6).

DNA damage activates kinase CK2, and then CK2 regulates mismatch repair, nucleotide excision repair (NER), non-homologous end joining (NHEJ), and homologous recombination (HR) through phosphorylating various DNA repair proteins (42). Silmitasertib (CX-4945) is a highly selective and potent CK2 inhibitor (43). Silmitasertib stabilized ATF4 when co-treated with PG3 or cisplatin (**Figure 6H**). As for PG3 and cisplatin treatment, silmitasertib rescued ATF4 degradation induced by PG3 and cisplatin. Importantly, the triple treatment of PG3 plus cisplatin and silmitasertib caused more potent apoptosis than PG3 and cisplatin, suggesting that stabilization of ATF4 is beneficial for increasing cytotoxicity against cancer cells (**Figure 6H**). Our data indicate that CK2 regulates ATF4 degradation induced by cisplatin in FaDu cells. In summary, a signaling pathway of ATF4 degradation is proposed based on the experiments (**Figure 6I**).

## Discussion

The ISR is considered a different target for cancer therapy distinct from targeting oncogenes (44). On one hand, as reported in our previous studies, ATF4 regulates expression of a subset of pro-apoptotic p53 target genes (*PUMA, NOXA, DR5, p21* etc.) and leads to apoptosis in *TP53*-mutated cancer cells (15, 16). On the other hand, in the ISR field, previously most of the attention for anti-cancer drug development was given to PERK, and PERK inhibitors such as GSK2606414 (45) and HC-5404(46), or PERK activator CCT020312 that inhibited triple-negative breast cancer and prostate cancer *in vitro* and *in vivo* by activating a PERK-ATF4-CHOP pathway without induction of ER stress (47, 48).

Recently, ISR and HRI were discovered to play an important role in cancer cell fitness and progression (44). Thus HRI becomes an attractive target for the development of anti-cancer drugs (44, 49, 50). HRI activator BTdCPU promotes eIF2α phosphorylation and induces apoptosis in dexamethasone-resistant multiple myeloma (49). Dihydroartemisinin activates HRI and leads to apoptosis through an HRI-ATF4-NOXA pathway (51, 52). Both BTdCPU and dihydroartemisinin lead to NOXA-mediated Mcl-1 degradation and inhibition (51, 52). HRI activation is involved in the innate immune response to bacterial infection (31) and upregulation of PD-L1 on cancer cell surface due to inhibition of heme synthesis, which causes ISR and selective translation of PD-L1 mRNA that has 3 uORFs in its 5’-UTR (53).

Mitochondrial stress induced by various mechanisms frequently leads to cancer cell apoptosis through activation of HRI and an HRI-eIF2α-ATF4 pathway (16, 33, 54–56). ONC201 and ONC212 bind and activate mitochondrial protease ClpP, leading to mitochondrial stress, which results in ISR through HRI (16, 56). CCCP (carbonyl cyanide m-chlorophenyl hydrazone) or oligomycin inhibits the electron transportation chain (ETC) and oxidative phosphorylation leads to mitochondrial stress and activation of ISR through HRI (54, 55).

DNA damage triggers the ISR and selective translation of DNA damage repair proteins and p21 (12, 13, 28). The functions of DNA damage-triggered ISR include but might not be limited to (1) quick translation of DDR proteins for DNA repair, (2) quick translation of p21 together with inhibition of global protein synthesis to further stop the cell cycle for DNA repair. HNSCC cells often exhibit elevated levels of ISR compared to normal cells. The ISR enables cancer cells to not only survive but also to enhance proliferation and progression within a stress-filled tumor microenvironment (TME) (57). Since intense or persistent ISR leads to cell death, we assume that the combination of ISR inducers plus DNA damaging drugs provide prolonged or intense ISR, leading to apoptosis in *TP53*-deficient cancer cells. We found that when *TP53*-mutated HNSCC cells were treated with PG3 and cisplatin, synergistic inhibitory effects, intense and persistent induction of ISR, and potent apoptosis were achieved (**Figure 1** and **Figure 2**).

To unravel the mechanism of driving enhanced therapeutic efficacy, through the ISR, we examined ISR signaling, focusing on eIF2α phosphorylation, ATF4, and NOXA. Addition of PG3 to cisplatin profoundly increased eIF2α phosphorylation compared to cisplatin or PG3 alone (**Figure 1F** and **G**).

We previously reported that PG3 activates HRI and leads to eIF2α phosphorylation (16). Therefore, we determined which eIF2α kinase was activated by cisplatin. Using CRISPR/Cas9 gene knockout and siRNA knockdown technologies, we uncovered that cisplatin activates HRI in FaDu cells (**Figure 5**). We infer that both PG3 and cisplatin small molecules converge on HRI activation, resulting in additive or synergistic phosphorylation of eIF2α.

However, it remains elusive how cisplatin activates HRI. Fu, *et al.* reported that mitochondrial DNA damage leads to the ISR and ATF4 activation through an ATAD3A-DELE1-HRI pathway (33). Hamada, *et al.* demonstrated that reactive oxygen species (ROS) lead to HRI activation (58). Cisplatin enters mitochondria and causes mitochondrial DNA damage (59), and cisplatin covalently binds to glutathione (GSH), leading to the production of ROS (3, 59). Hence, HRI activation by cisplatin is possibly due to either cisplatin-induced mtDNA damage or the production of ROS in cells.

We found that ATF4 modulates both cisplatin-or PG3 and cisplatin-induced NOXA upregulation, and ATF3 does not participate in the regulation of NOXA expression in *TP53*-mutated FaDu and Cal27 HNSCC cells (**Figure 4**). Sharma, *et al.* reported that ATF4 and ATF3 regulate cisplatin-induced upregulation of NOXA in *TP53*-deleted or -truncated HN8 and HN12 cells. Possibly the difference originates from different cell lines and the cisplatin concentration used. Another unexplained observation is that PG3 leads to PUMA-dependent apoptosis in *TP53*-mutated colorectal cancer cells (16). However, the combined treatment of PG3 and cisplatin causes NOXA-dependent apoptosis in HNSCC cells. As mentioned previously, NOXA and PUMA compete to bind to Mcl-1. Silencing PUMA led to upregulation of NOXA and increased PARP cleavage and apoptosis induced by cisplatin or PG3 and cisplatin (**Figure 3A, B**, and **C**). On the other hand, although knockdown of *NOXA* induced upregulation of PUMA, it did not increase PARP-cleavage; instead, it inhibited PARP-cleavage induced by cisplatin or PG3 and cisplatin, suggesting PUMA does not contribute to the apoptosis under the experimental conditions (**Figure 3A, B,** and **C**). Therefore, either NOXA dominates the competition and leads to cellular apoptosis, or PUMA is not required for PG3 and cisplatin-induced apoptosis. In summary, PG3, as an ISR inducer, increases the efficacy of cisplatin treatment by enhancing the ISR and NOXA-mediated apoptosis.

β-TrCP1 is activated by DNA damage via activation of NAE and plays an important role in DNA damage repair (40). As previously mentioned, DNA damage activates CK2, and CK2 plays a crucial role in DNA repair. However, the exact mechanisms of CK2 activation are still unknown (42). DNA damage activates kinase CK1δ through ATM-mediated phosphorylation of CK1δ, participating in cell cycle regulation and DNA repair (41, 60). Pevonedistat inhibits β-TrCP1 through inhibition of NAE. Pevonedistat received a Breakthrough Therapy Designation from the FDA for the treatment of higher-risk myelodysplastic syndrome (HR-MDS) in 2020. Pevonedistat has been investigated in combination therapies for the treatment of blood cancers (NCT04266795, NCT03330821) or solid tumors, such as lung, liver, and cholangiocarcinoma (NCT03965689, NCT04175912). Silmitasertib (CK2 inhibitor) was granted orphan drug status by the U.S. Food and Drug Administration for cholangiocarcinoma in 2017 (61). Currently, it is being investigated in combination with irinotecan, temozolomide, and vincristine for the treatment of relapsed or refractory solid tumors, such as neuroblastoma, Ewing’s sarcoma, and osteosarcoma (NCT06541262). CK1δ inhibitors have not entered clinical trials as of the writing of this manuscript. However, CK1δ is a promising target for breast cancer therapeutics, and a selective inhibitor, SR-3029, has shown efficacy against breast cancer subtypes overexpressing CK1δ (62).

Degradation of ATF4 was reported as an acquired-resistance mechanism to cisplatin treatment and persistent ISR activation (6, 63), and the stabilization of ATF4 restores the sensitivity to cisplatin and ISR (6, 63). We noted a decrease in ATF4 levels in Cal27 and FaDu HNSCC cells after treatments with DNA damaging cisplatin, CPT (**Figure 6A** and **B**), or PG3 and cisplatin (**Figure 1F, G, 3G,** and **6G**). This suggests that the combined treatment of PG3 and cisplatin might result in acquired resistance via degradation of ATF4. To resolve this, we targeted the degradation mechanism of ATF4 by inhibiting β-TrCP1, CK1δ, or CK2, respectively. Each approach successfully blocked ATF4 degradation induced by cisplatin or PG3 and cisplatin and enhanced apoptosis (**Figure 6**). It remains unclear why DNA damage causes eIF2α phosphorylation and induction of the ISR but leads to degradation of ATF4. Our results provide a rational strategy of triple treatments, such as ISR inducer plus DNA damaging drug plus β-TrCP1 inhibitor/CK1δ inhibitor/CK2 inhibitor, to achieve potent and prolonged anti-tumor effects to overcome chemoresistance.

## Materials and Methods

### Cell lines and reagents

#### *TP53*-mutant cell lines

FaDu (R248L), Cal27 (H193L), PANC-1(R273H), HT29 (R273H), and normal fibroblast cell lines HFF-1, and IMR90 cells were purchased from ATCC. Cells were routinely checked for mycoplasma, and all cell lines underwent STR authentication. HT29 was cultured in McCoy’s 5A medium (Gibco) with 10% fetal bovine serum (FBS, Invitrogen). FaDu, Cal27, PANC-1, HFF-1, and IMR90 were cultured in Dulbecco’s modified Eagle’s medium (DMEM, Gibco) with 10% fetal bovine serum (FBS, Invitrogen) at 37°C in humidified air with 5% CO_2_.

#### Chemicals

PG3 (Provid Pharmaceuticals, New Jersey). Cisplatin, 2Bact, silmitasertib, and pevonedistat (MedChemExpress); Z-VAD-FMK, MG-132, and SB216763 (Selleckchem); Thapsigargin (Tocris Bioscience); D4476 (Cayman Chemicals), and ONC201 (Chimerix, Durham, NC).

#### Cell viability assay

Cells were seeded into 96-well black plates (6 x 10^3^ cells/well). The cells were treated with different concentrations of compounds or dimethyl sulfoxide (DMSO) as a control for 72 hours. The cell viability was assessed by CellTiterGlo bioluminescent cell viability assay (Promega), following the manufacturer’s protocol. Bioluminescence imaging was measured using the IVIS imager. The percentage of cell viability and synergy scores was calculated against the respective DMSO control using the software Combenefit using the model HAS and displayed as surface plots where dose-responses were integrated (17).

#### Western blotting

After treatment, protein lysates were collected for Western blot analysis. A total of 15 μg of protein was used for SDS-PAGE. After primary and secondary antibody incubations, the signal was detected by a chemiluminescence detection kit, imaged by Syngene (Imgen Technologies). Antibodies for PUMA, CK1δ, and HRI were from Santa Cruz Biotechnology; for caspase-8, cleaved caspase-8, caspase-9, caspase-3, cleaved-caspase-3, PARP, cleaved-PARP, GFP, CDK2, p-CDK2(T160), Mcl-1, p-Mcl-1(S64), β-TrCP1, PERK, PKR, GCN2, eIF2α, p-eIF2α (Ser51), ATF3, and ATF4 were from Cell Signaling Technology. NOXA was from Milipore Sigma. CReP was from Proteintech. Ran was from BD Biosciences.

#### siRNA knockdown

siRNA silencing experiments were performed by transfecting either 80 pmoles of indicated siRNA(s), or scramble siRNA using RNAiMAX (Invitrogen), following the manufacturer’s protocol. Transfected cells were treated with drugs 24-or 48-hours post-transfection. The control siRNA and siRNAs for human NOXA, PUMA, CK1δ, and HRI were purchased from Santa Cruz Biotechnology. siRNAs for human ATF4, ATF3, and PERK were from ThermoFisher Scientific.

#### Production of PANC1 and FaDu knockout cells

Lentivirus was produced by transfecting Lenti-X-293T cells (Takara #632180) with the following plasmids: transfer plasmid, pRSV-Rev (Addgene #12253), pMDLg/pRRE (Addgene #12251), and pMD2.G (Addgene #12259) using Lipofectamine 2000 (ThermoFisher 11668019) per the manufacturer’s protocol. The transfer plasmid used are as follows: lentiCas9-Blast (Addgene #52962), non-targeting control gRNA (Addgene #80263), HRI g2 (Addgene #77050), PKR g1 (Addgene #75637), PERK g1 (Addgene #77166), PERK g2 (Addgene #77168), GCN2 g1 (Addgene #75877), and GCN2 g2 (Addgene #75876).

To produce the PANC1 knockout cells, PANC1 was transduced with lentivirus containing the lentiCas9-Blast transfer plasmid and selected with blasticidin to produce PANC1-Cas9 cells. After blasticidin selection was finished, cells were then transduced with either lentivirus containing transfer plasmids for the non-targeting control gRNA or the indicated targeting gRNA. They were then selected with puromycin. Two guide NOXA RNAs were designed using Integrated DNA Technologies (Coralville, Iowa 52241) guide RNA design tool. Control sgRNA and the two sgNOXAs were synthesized by GenScript (Piscataway, NJ). sgRNAs were transfected into FaDu cells using Lipofectamine CRISPRMAX (Invitrogen) following the manufacturer’s protocol.

#### Flow cytometry

Cell Death Analysis —Zombie Violet staining and flow cytometry were used to determine the degree of cellular death. Cells were seeded at 5 ×10^5^ cells/well in 6-well plates. Cal27 cells were treated with PG3, cisplatin, 2Bact, PG3 plus 2Bact, cisplatin plus 2Bact, PG plus cisplatin, and PG3 plus cisplatin, respectively for 12 hours. The cells were harvested and stained with Zombie Violet dye following the manufacturer’s protocol (BioLegend). Then, flow cytometry analysis was performed according to the manufacturer’s protocol (CytoFlex, Beckman Coulter, Brea, California).

#### Production of ATF4 reporter cell lines

Lentivirus was produced by transfecting Lenti-X-293T cells (Takara #632180) with the following plasmids: pSMALB-ATF4.5 (Addgene #155032), pRSV-Rev (Addgene #12253), pMDLg/pRRE (Addgene #12251), and pMD2.G (Addgene #12259) using Lipofectamine 2000 (ThermoFisher 11668019) per the manufacturer’s protocol. 16-24 hours of post-transfection, the media was changed to fresh DMEM. At 48-72 hours post-transfection, the media was collected, passed through a 0.45 micrometer PES syringe filter (Millipore SLHPR33RS) before being used to transduce the cells. FaDu and Cal27 were then transduced with lentivirus produced as above for 24 hours. After cells were expanded, they were then sorted for tagBFP positive cells.

#### Live cell imaging

Cal27-ATF4-GFP cells were plated at a density of 10,000 cells per well of a 96-well plate in FluoroBrite® DMEM imaging medium containing 10% FBS and 4 mM glutamine (Gibco). The cells were treated with thapsigargin (0.2 μM) and 10 μM ONC201, respectively, for 24 hours. Imaging was performed using the ImageXpress® Micro Confocal High-Content Imaging system (ImageXpress, Inc.).

#### Statistical analysis

All results were obtained from triplicate experiments, unless otherwise indicated. Statistical analyses were performed using Prism10 software (GraphPad Software, Inc.) and the student t test. Statistical significance was determined by P < 0.05.

## Author contributions

X.T., and W.S.E-D. conceptualized the project and all experiments that were performed. X.T. and P. R.S. were involved with the technical performance of all experiments. X.T., P.R.S., and W.S.E-D. were involved in all the data analysis and discussion of the results.

## Acknowledgements

W.S.E-D. is an American Cancer Society Research Professor and is supported by the Mencoff Family University Professorship at Brown University. This work was presented in part at the 2025 annual meeting of the American Association for Cancer Research.

pSMALB-ATF4.5 was a gift from John Dick & Peter van Galen (Addgene plasmid # 155032)(18). pRSV-Rev (Addgene plasmid # 12253), pMDLg/pRRE (Addgene plasmid # 12251), and pMD2.G Addgene plasmid # 12259) were gifts from Didier Trono(64). lentiCas9-Blast was a gift from Feng Zhang (Addgene plasmid # 52962)(65). non-targeting control gRNA (BRDN0001148862, Addgene plasmid # 80263), EIF2AK1 gRNA (BRDN0001162524, Addgene plasmid # 77050), EIF2AK2 gRNA (BRDN0001146342, Addgene plasmid # 75637). EIF2AK3 gRNA (BRDN0001145848, Addgene plasmid # 77166), EIF2AK3 gRNA (BRDN0001146966, Addgene plasmid # 77168). EIF2AK4 gRNA (BRDN0001147521, Addgene plasmid # 75877), and EIF2AK4 gRNA (BRDN0001146469, Addgene plasmid # 75876) were gifts from John Doench & David Root(66).

---

**Supplementary Figure 1.**
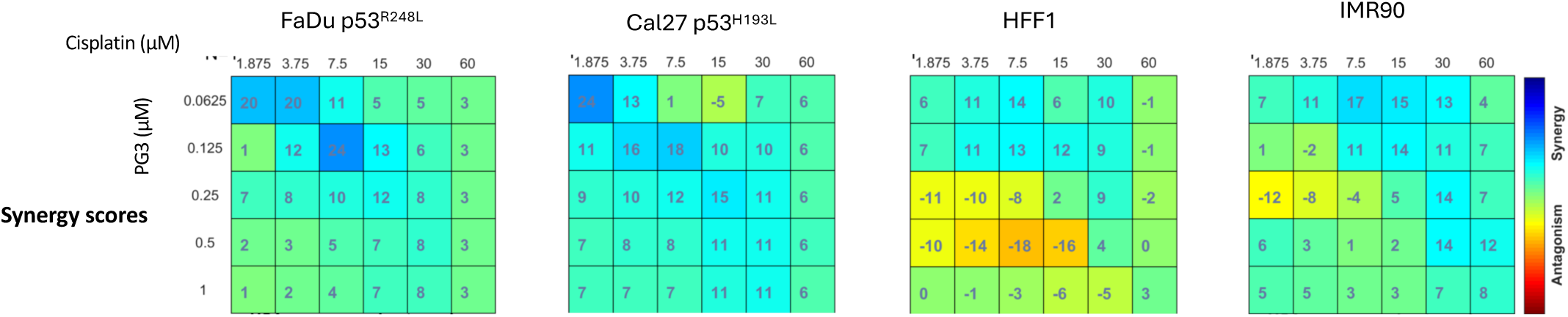
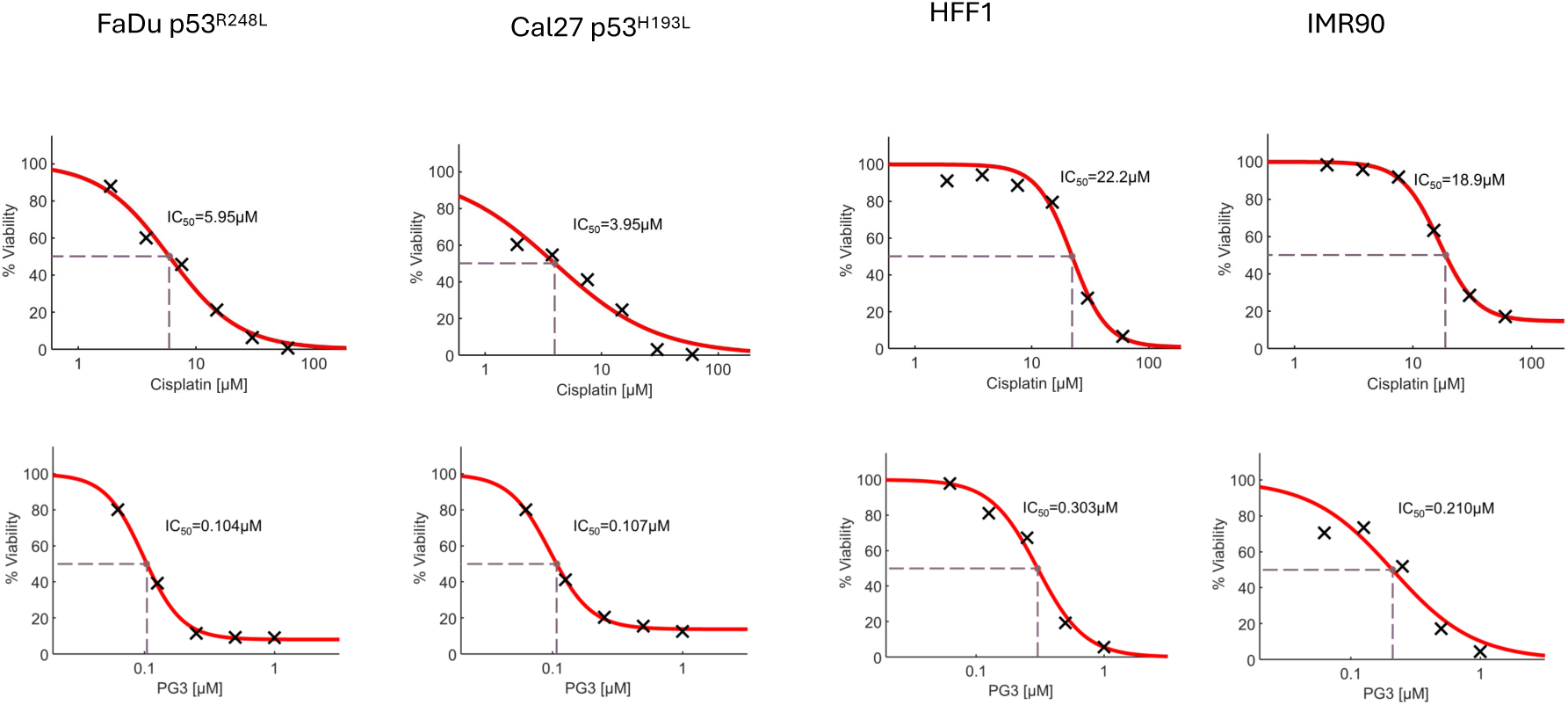
PG3 and cisplatin synergistically inhibit cell viability. Cell viability was measured in FaDu, Cal27, HFF-1, and IMR90 cells. Cells were treated with different concentrations of PG3, cisplatin, or DMSO for 72 h. Luciferase activity was imaged using the IVIS Imaging System after treatment. Cell viability data were normalized to those of DMSO treatment control in each cell line, and data analyses were performed using Combenefit software.

**Supplementary Figure 2.**
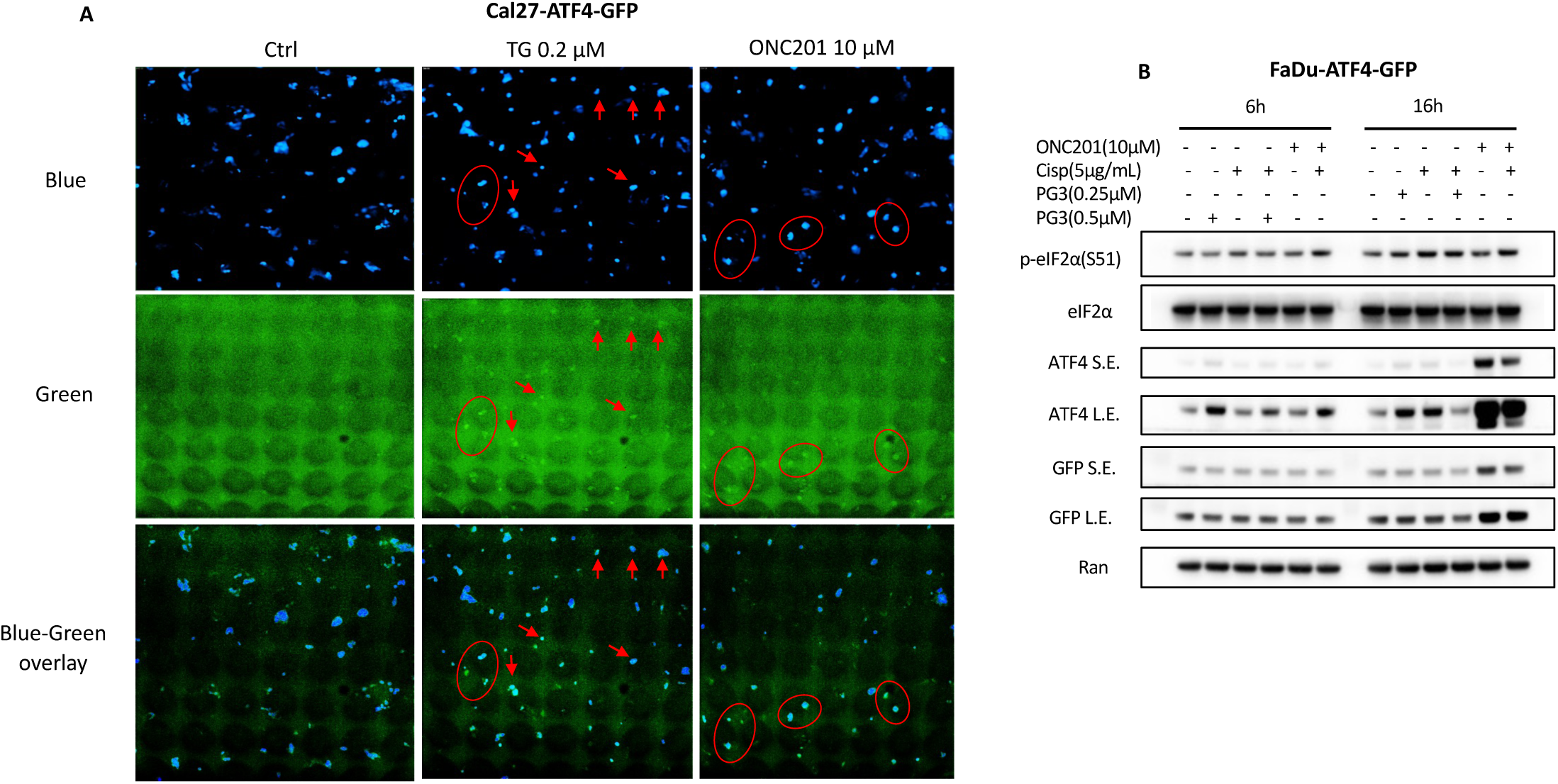
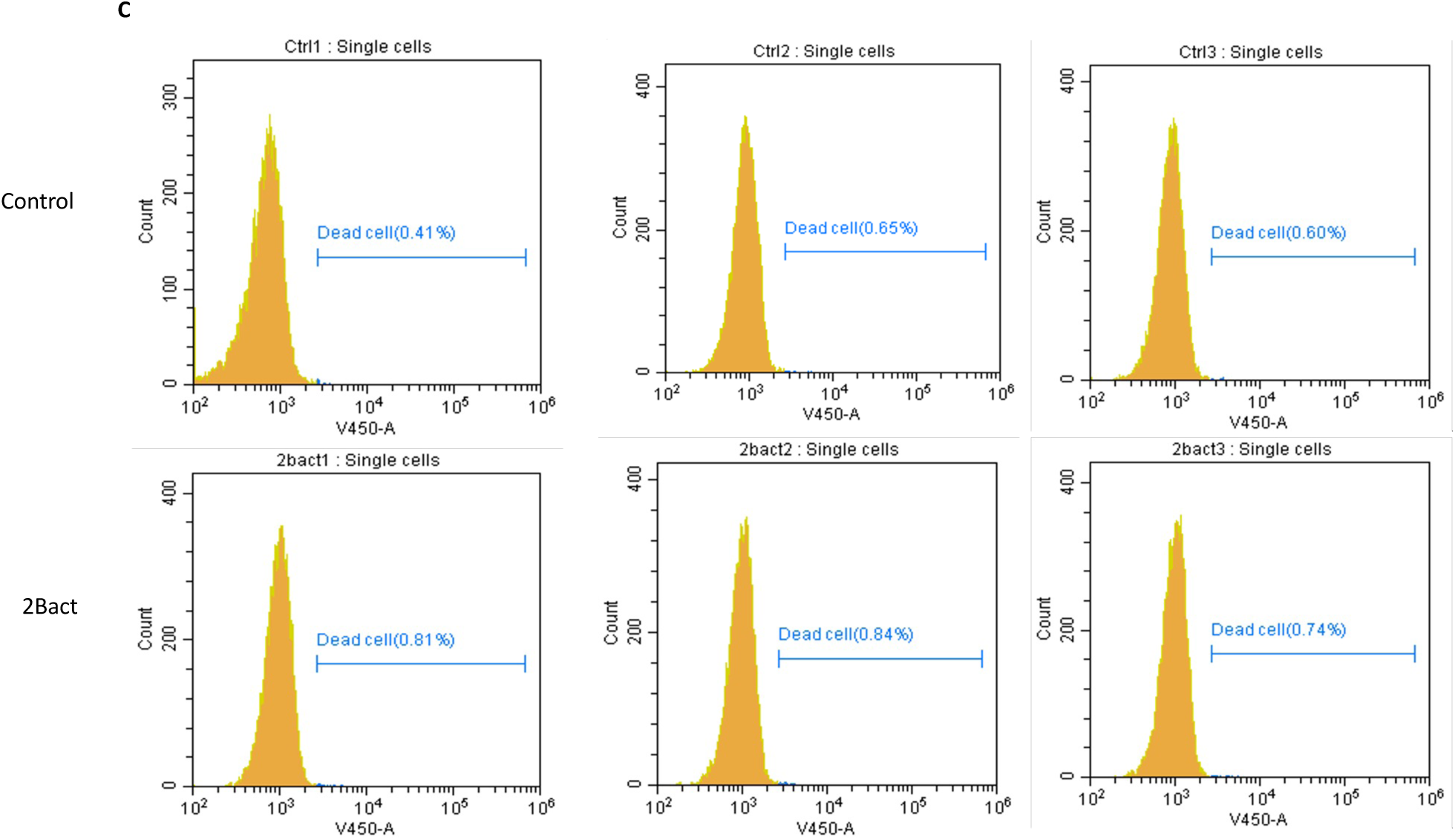
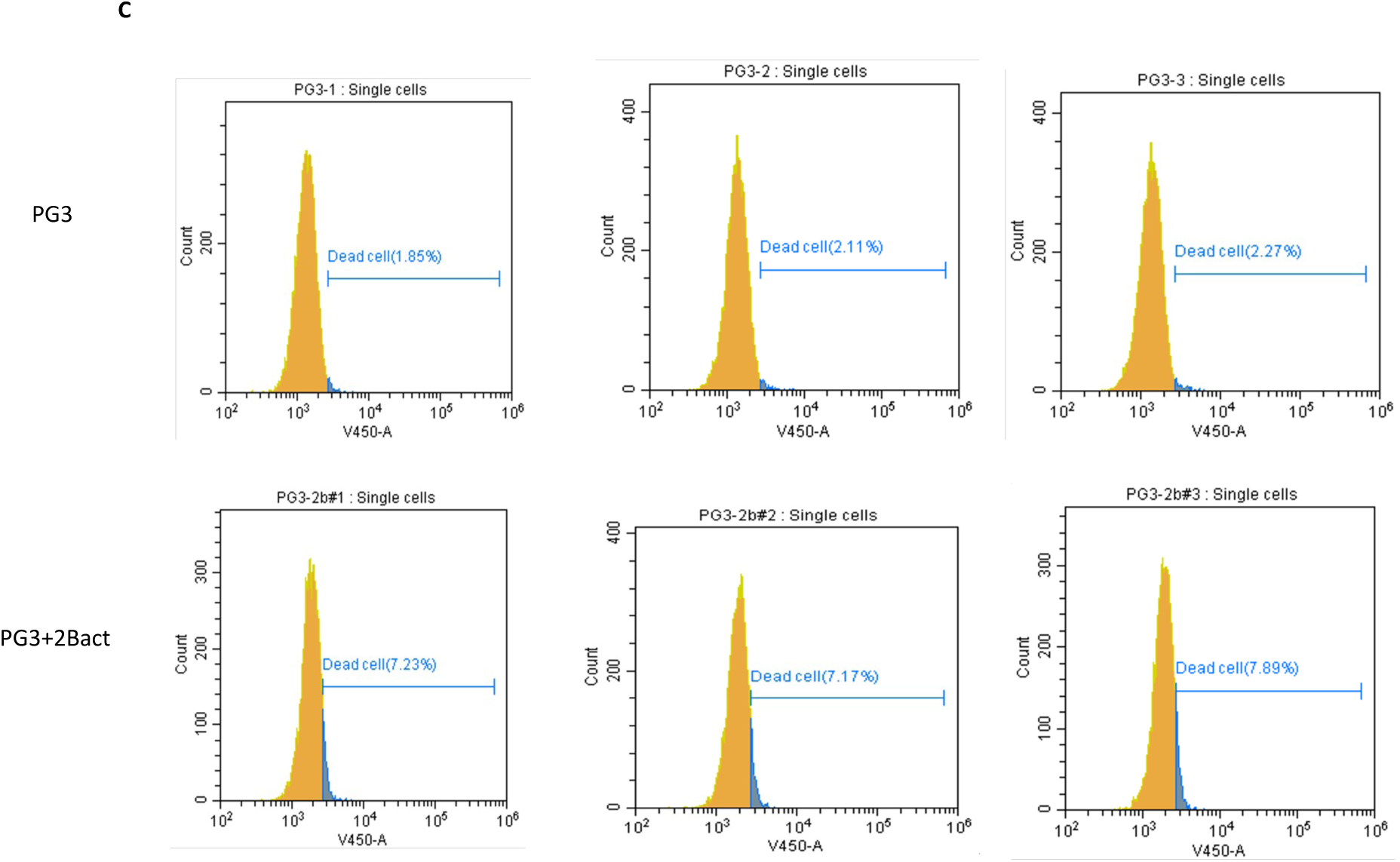
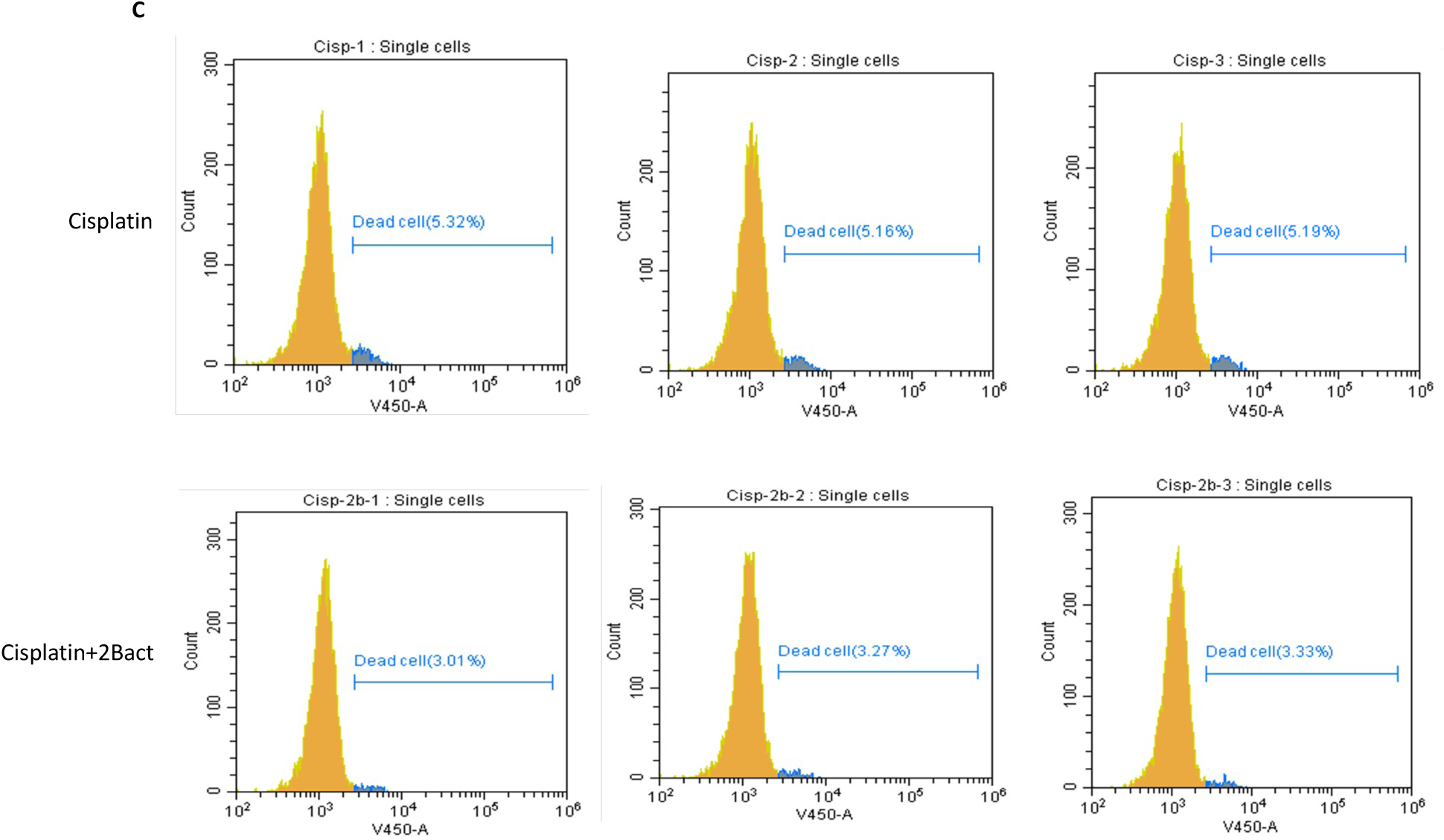
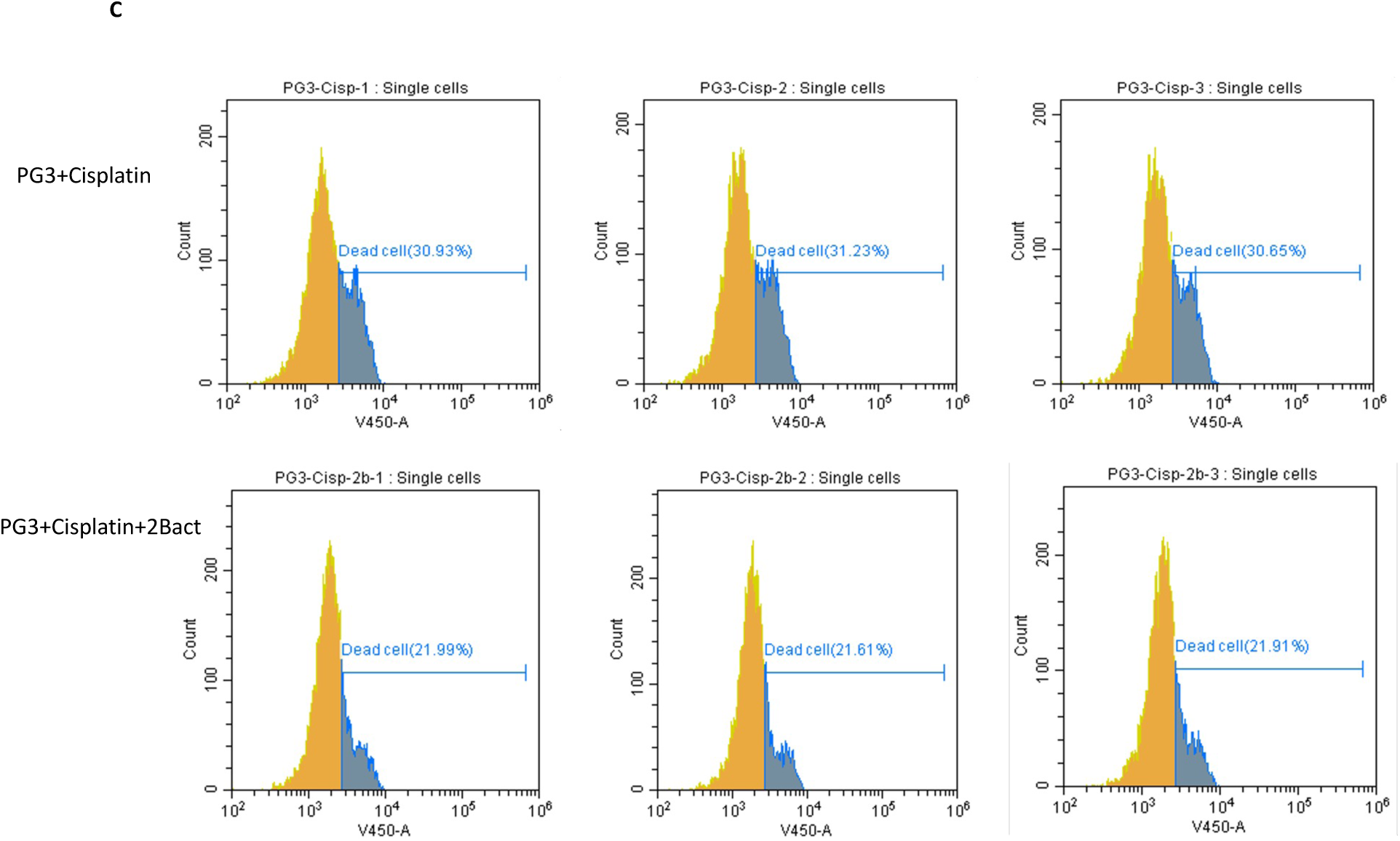
The ISR mediates cell apoptosis. (A) Cal27-ATF4-GFP reporter cells were treated with TG or ONC201 for 24 hours. Images were taken using the ImageXpress® Micro Confocal High-Content Imaging system. (B) FaDu-ATF4-GFP reporter cells were treated with PG3, cisplatin, and thapsigargin (TG) for 6 or 16 hours, respectively. Western blot was performed using the indicated antibodies.

